# Cortical inhibitory potentiation reverses maladaptive amygdala plasticity after noise-induced hearing loss

**DOI:** 10.64898/2026.04.02.716147

**Authors:** Bshara Awwad, Daniel B Polley

## Abstract

Across sensory modalities, peripheral deafferentation can induce maladaptive central plasticity that amplifies sensory responses while distorting affective processing, rendering innocuous stimuli unpleasant, intrusive, or even painful. Here, we examined the affective dimension of compensatory plasticity in a mouse model of hyperacusis by introducing noise-induced focal cochlear lesions and tracking maladaptive plasticity in the lateral amygdala (LA), a key limbic hub for sensory-affective valuation. Whereas control mice habituated to neutral sounds, mice with noise-induced hearing loss (NIHL) showed sustained LA hyperresponsivity and abnormally strong temporal coupling between calcium transients and pupil dilations, an autonomic index of arousal. NIHL also disrupted discriminative auditory threat learning, producing poorly selective, non-extinguishing enhancement of LA responses and freezing to both threatening and non-threatening sounds. We posited that potentiating inhibition in the higher-order auditory cortex (HO-AC), a prominent source of auditory sensory input to the LA, could reinstate normal affective sound processing after NIHL. Brief bouts of 40-Hz optogenetic activation of HO-AC parvalbumin-expressing inhibitory neurons (PVNs) durably reversed LA sensitization, normalized pupil-indexed arousal dynamics, and reinstated discriminative auditory threat memory. These findings identify cortical inhibitory potentiation as a strategy for reversing neural, autonomic, and behavioral signatures of distorted affective sound processing after peripheral sensory injury.

## INTRODUCTION

Peripheral sensory injuries produce a paradoxical combination of sensory loss and perceptual excess. This perceptual excess commonly takes three forms: phantom percepts in the absence of a physical stimulus, increased gain that amplifies the intensity of external stimuli, and distorted affective processing that renders innocuous stimuli unpleasant, intrusive, or even painful. For example, after traumatic limb injury, the loss of somatosensory afferent input can be accompanied by vivid pain and discomfort arising from a phantom limb that feels contorted in an unnatural position. Likewise, reduced afferent input from olfactory receptor neurons can occur alongside the perception of phantom odors (phantosmia) and a distorted perception of neutral odorants such that they are experienced as foul, burnt, or otherwise disgusting (parosmia). This combination of sensory loss and perceptual excess is especially prevalent in the auditory system, where approximately 15% of adults report hearing difficulties, 12% report the perception of phantom sounds (tinnitus), and 9% report that sounds are not only uncomfortably loud, but also unpleasant and aversive (hyperacusis)^1–3^.

Whereas maladaptive plasticity processes that give rise to the sensory dimensions of tinnitus and hyperacusis have been studied extensively, the neural basis of their affective dimension remains poorly understood. Neuroimaging studies have identified abnormally strong functional coupling between auditory cortex and limbic regions, including the amygdala^4–6^. Some models explain this coupling through disrupted gating mechanisms involving prefrontal cortex, ventral striatum, and thalamic reticular nucleus ^7–9^. We instead considered a more parsimonious mechanism rooted in excess central gain, a maladaptive compensatory plasticity triggered by hearing loss.

Central auditory circuits compensate for the loss of cochlear afferent input through disinhibition^10–15^. Following noise-induced hearing loss (NIHL), parvalbumin-expressing inhibitory neurons (PVNs) in the auditory cortex provide weaker feedforward inhibition to excitatory pyramidal neurons^16,17^, producing hyperactive network activity that is necessary and sufficient to account for loudness hypersensitivity^18^.

Deep-layer auditory cortical neurons that project to telencephalic and subcerebral targets exhibit especially pronounced hypersensitivity and hyperresponsivity after NIHL^19,20^. The lateral amygdala (LA), positioned on the dorsal surface of the basolateral amygdala complex, serves as a principal gateway through which auditory inputs engage limbic circuits^21–23^. LA excitatory neurons receive dense monosynaptic input from glutamatergic corticoamygdalar projection neurons in higher-order auditory cortex (HO-AC), as well as from higher-order auditory and multisensory thalamic nuclei^24,25^. Associative plasticity within the LA and its presynaptic auditory inputs imbues neutral sounds with affective significance by supporting selective memories of threat-predicting sounds and engaging downstream regulators of defensive behavior, such as freezing, and autonomic arousal, such as pupil dilation^26–32^. A similar combination of reduced motor reactivity and enhanced pupil dilation has been reported in individuals with bothersome tinnitus and hyperacusis in response to negative valence sounds, raising the possibility that excess central auditory gain after hearing loss and disrupted affective processing of arousal and valence may be linked^33^.

Here, we posited that distorted affective sound processing after NIHL reflects the conjunction of two plasticity processes: disinhibition and excess cortical gain in response to hearing loss, and a resulting maladaptive engagement of LA circuits that assign affective valence and arousal to auditory stimuli. To test this idea, we performed chronic fiber photometry of LA excitatory neurons before and after NIHL and related sound-evoked LA activity to autonomic indices of arousal and discriminative auditory threat learning. We also tested a corollary prediction: if reduced PVN-mediated inhibition in auditory cortex is a critical failure point that drives excess gain in cortical projection neurons and downstream dysregulation of auditory threat circuits, then potentiating cortical PVN inhibition should reinstate normal downstream affective processing after NIHL. In recent work, we found that a single bout of optogenetic activation of auditory cortical PVNs durably reversed behavioral loudness hypersensitivity after NIHL^18^. Here, we show that the same intervention also reverses distorted affective processing associated with NIHL.

## RESULTS

### Lateral amygdala sensitization – rather than habituation – to familiar sounds after noise-induced hearing loss

Noise-induced cochlear damage reduces afferent input to central auditory pathways and produces auditory hyperresponsivity in auditory cortical projection neurons^19,20,34–37^. We hypothesized that hyperresponsivity in long-range projection neurons could disrupt affective sound processing in postsynaptic limbic brain areas such as the lateral amygdala (LA) and downstream circuits that regulate autonomic responses to sound (**Figure 1a**). To test this, we exposed mice to 2 hours of octave-band noise that induced focal sensorineural damage in the high-frequency base of the cochlea (**Figure 1b**). After noise-induced hearing loss (NIHL), auditory brainstem responses (ABR) evoked by sound frequencies > 16kHz were absent, while responses to frequencies below 16 kHz were spared (**Figure 1c**, statistics reported in the figure legends throughout). To document changes in LA and autonomic sound processing after NIHL or an innocuous sham exposure, we expressed GCaMP6s in LA excitatory neurons and performed fiber-based bulk calcium imaging and pupil measurements in head-fixed mice over a 7-day period before and after NIHL and sham exposure (**Figure 1d-e**). Post-mortem reconstructions of fiber tip placement and LA boundaries are provided in **Supplemental Figure 1**.

**Figure 1.**
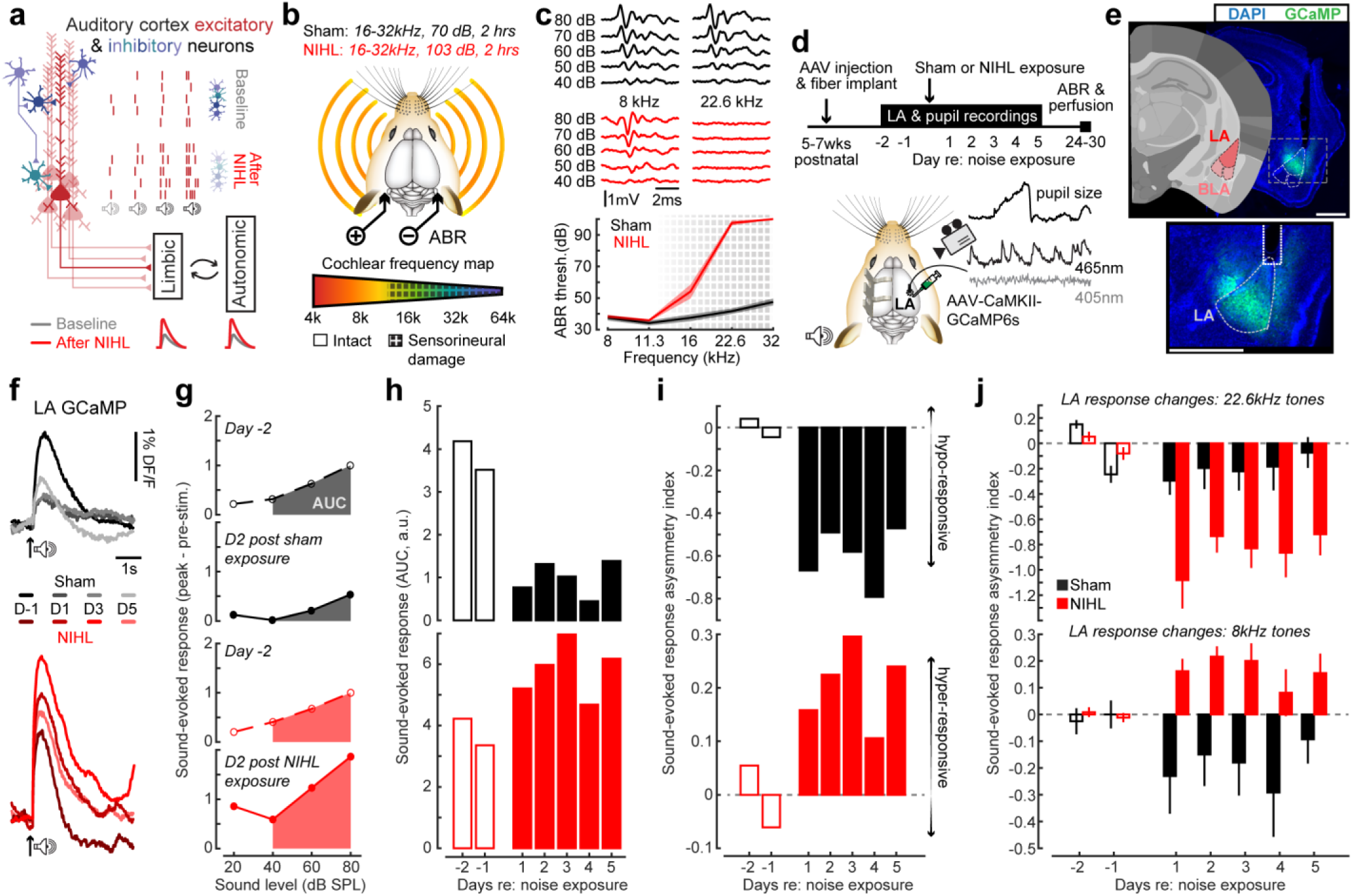
Habituation and sensitization of sound-evoked lateral amygdala responses. (a) A schematic model for abnormal limbic and autonomic recruitment following NIHL. Cartoon illustrates AC excitatory projection neurons (red) and inhibitory interneurons (blue shades). Reduced afferent input triggers a compensatory reduction in local inhibition, causing a steeper growth in spiking with sounds of increasing intensity (i.e., excess central gain, the purported generator of loudness sensitivity). Hyperactive AC projection neurons directly innervate limbic brain regions like the LA, potentially introducing knock-on hyperresponsivity and enhanced recruitment of downstream brain centers that regulate the autonomic nervous system. (b) Cartoon illustrates the noise exposure protocols and the region of cochlear sensorineural damage after NIHL. Mice were exposed to 16–32 kHz octave-band noise for 2 hours at 70 dB SPL (sham) or 103 dB SPL (NIHL). (c) *Top:* ABR waveforms from example sham and NIHL mice elicited by 8 kHz and 22.6 kHz tones. *Bottom*: mean ± SEM ABR threshold for sham (N = 6) and NIHL (N = 6) mice. NIHL mice showed elevated thresholds restricted to high frequencies (LME, Frequency × Group interaction, F(4,48) = 124.69, p = 1.0×10⁻²⁴; post-hoc pairwise comparisons: 8 kHz: p = 0.73; 11.3 kHz: p = 0.73; 16 kHz: p = 0.028; 22.6 kHz: p = 1.84×10⁻⁹; 32 kHz: p = 1.45×10⁻⁹). Scale bars, 1 μV and 2 ms. (d) *Top*: Experimental timeline. *Bottom*: AAV-CaMKII-GCaMP6s was injected into the right LA and a fiber optic cannula was implanted. Fiber-based bulk fluorescence measurements were made in the LA at the GCaMP excitation wavelength (465nm) and a calcium-insensitive control wavelength (405nm). Pupil dynamics were measured simultaneously. (e) *Top:* Coronal sections depict the caudal portion of the LA targeted for injection. Nomenclature and reference image are adapted from the Allen Institute for Brain Science. Scale bar = 1mm. *Bottom:* Fluorescence micrographs show GCaMP6s expression and estimated position of the optic fiber (dotted white line), and approximate boundaries of the LA (dashed white line). Scale bar = 0.5mm. See also Figure S1. (f) Sound-evoked GCaMP responses before (D-1) and after noise exposure (D1, D3, D5) in an example sham (black) and NIHL (red) mouse. Responses were elicited by an 8kHz tone at 80 dB SPL. Scale bars = 1% ΔF/F and 1 s. (g) Sound level LA growth functions from the same example mice before (dashed line) and after (solid line) noise exposure. Shaded area depicts the area under the curve (AUC) from 40-80 dB SPL, the metric used throughout to capture the sound-evoked LA response for each recording session. (h) Sound-evoked AUCs for the same two example mice across the two baseline (open bars) and five post-exposure (solid bars) recording sessions. (i) Changes in sound-evoked AUC are shown for the same two example mice with an asymmetry index ([Session X – Baseline_mean_] / [Session X + Baseline_mean_]) where negative values indicate hypo-responsiveness (habituation) and positive values indicate hyper-responsiveness (sensitization). (j) Mean ± SEM sound-evoked response asymmetry index across days relative to noise exposure for 22.6 kHz (top) and 8 kHz (bottom) tones (N = 6/6, sham/NIHL). LA responses habituated to 8 and 22.6 kHz tones in sham mice, while NIHL mice exhibited suppressed responses to 22.6 tones that correspond to damaged regions of the cochlea and enhanced responses to 8kHz tones that activated spared regions of the cochlea (Linear mixed effect model with frequency and session as repeated measures and group as a between-subjects factor; frequency × session × group interaction, F = 6.89, p = 2.45 × 10⁻⁶).

When mice were first introduced to neutral sounds like pure tones, LA excitatory neurons exhibited robust calcium responses that increased monotonically with sound level (**Figure 1f-g**). LA responses diminished throughout the week in controls, as mice became habituated to the auditory stimuli and test protocol^22^ (**Figure 1h-I, top**). By contrast, LA responses to 8kHz tone bursts were increased far above baseline levels after NIHL, reflecting an increased gain in neural responses to spared sound frequencies that mirrors many prior descriptions of enhanced AC response gain (**Figure 1h-I, bottom**)^18,20,34,38–40^. This shift from habituation to sensitization was only observed for spared low-frequency stimuli, as LA responses to high-frequency tones that activate cochlear regions with sensorineural damage were significantly reduced after NIHL (**Figure 1j**).

### Autonomic hyper-arousal and hyper-correlation with amygdala activity

The amygdala coordinates recruitment of the sympathetic nervous system in response to salient environmental stimuli^41^. To determine whether LA habituation or sensitization was accompanied by commensurate changes in pupil-indexed arousal, we measured sound-evoked pupil dilations during the same 8kHz tone trials that elicited habituated or sensitized LA responses. In control mice, pupil dilations evoked by 8kHz tones were only observed at the highest sound level and were relatively stable across the recording period (**Figure 2a and 2b**). Following NIHL, tone-evoked pupil dilations were larger in amplitude and more sensitive to moderate sound levels (**Figure 2b and 2c**).

**Figure 2.**
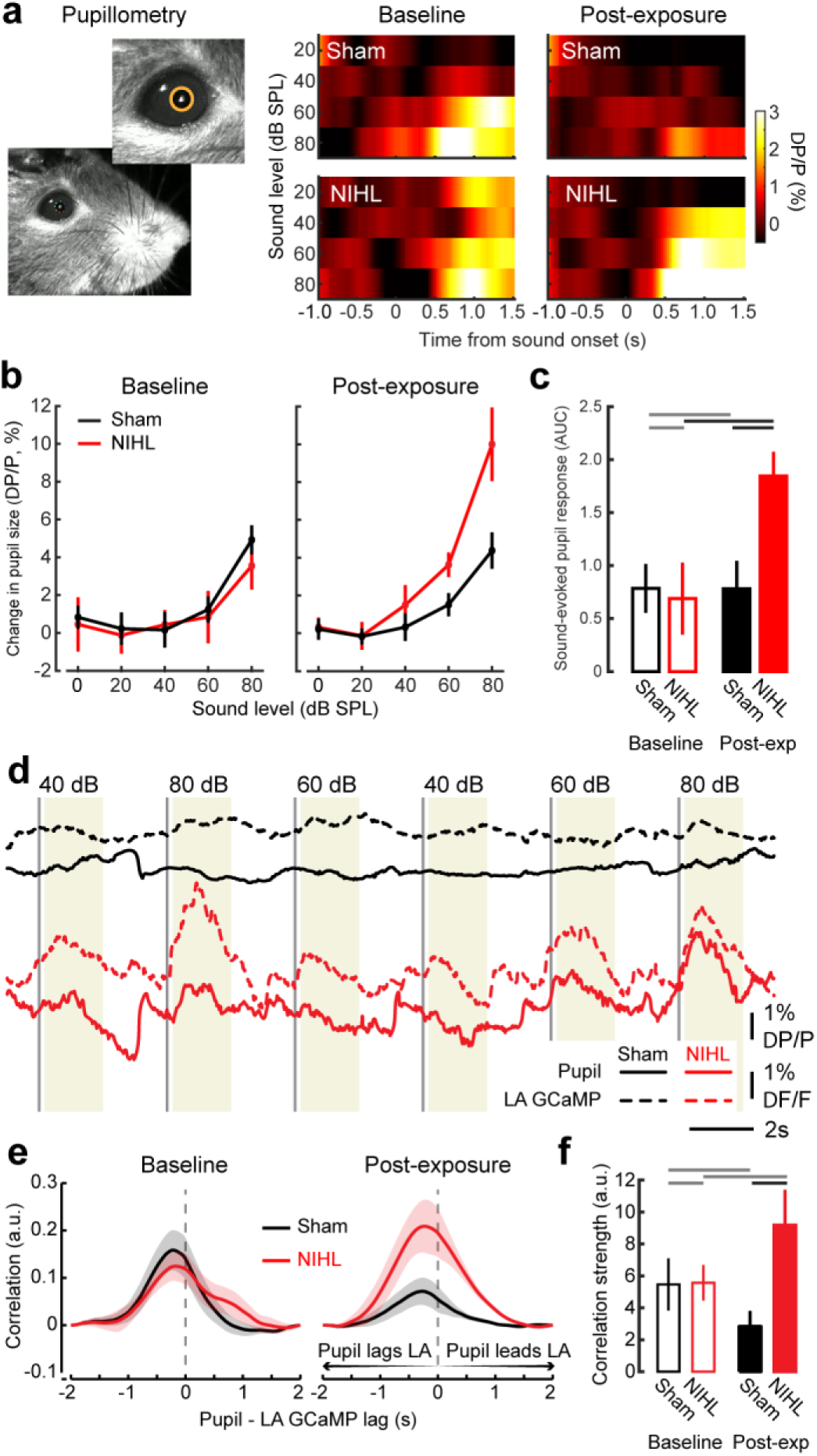
Enhanced pupil-indexed arousal and LA–pupil coupling after noise-induced hearing loss. (a) Pupillometry was performed concurrently with LA fiber photometry recordings. Heatmaps depict mean sound-evoked pupil dilation by 8kHz tones (ΔP/P, %) as a function of sound level for a representative sham (*top)* and NIHL (*bottom*) mouse across baseline and post-exposure sessions. (b) Mean ± SEM sound-evoked change in pupil size as a function of sound level at baseline and post-exposure for sham (N = 6) and NIHL (N = 6) mice. (c) Sound-evoked pupil response (AUC) at baseline and post-exposure. Sound-evoked pupil dilation was stable in sham-exposed mice but significantly enhanced after NIHL (LME with session as a repeated measure and group as a between-subjects factor; session × group interaction, F = 7.59, p = 0.013). Black and gray horizontal bars indicate significant and non-significant differences, respectively, based on pairwise comparisons with Holm–Bonferroni correction: NIHL baseline vs. post-exposure, p = 0.006; sham baseline vs. post-exposure, p = 0.99; sham vs. NIHL baseline, p = 0.78; sham vs. NIHL post-exposure, p = 0.018. Error bars, SEM. (d) Simultaneous LA GCaMP (dashed) and pupil (solid) traces from representative sham and NIHL mice during presentation of tones at varying levels (dB SPL labels above). Vertical dashed lines indicate stimulus onset. Yellow shading denotes the response analysis window. Scale bars, 1% ΔF/F, 1% ΔP/P, and 2 s. (e) Mean ± SEM normalized cross-correlation between LA GCaMP and pupil signals at baseline (*left*) and post-exposure (*right*) for sham (black, N = 6) and NIHL (red, N = 5) mice. Negative lags indicate that the pupil signal lags behind LA activity. LA–pupil coupling was enhanced in NIHL mice after noise exposure, with the pupil signal lagging LA activity. (f) LA-pupil temporal coupling, quantified as the AUC of positive cross-correlation values, at baseline and post-exposure. LA–pupil coupling was selectively enhanced in NIHL mice after noise exposure (LME with session as a repeated measure and group as a between-subjects factor; session × group interaction, F = 7.27, p = 0.015). Pairwise comparisons with Holm–Bonferroni correction: sham vs. NIHL baseline, p = 0.80; sham vs. NIHL post-exposure, p = 0.019; NIHL baseline vs. post-exposure, p = 0.11; sham baseline vs. post-exposure, p = 0.28. Error bars, SEM.

The LA lies upstream of brainstem circuits that regulate sympathetic effectors such as the pupil. However, LA activity dynamics could be tightly coupled with pupil dilations in conditions where sensory inputs were highly salient. Indeed, we noted a close temporal correspondence between sound-evoked LA responses and pupil dilations after NIHL that was not observed in control mice (**Figure 2d**). To quantify changes in temporal correlation independent of the aforementioned changes in LA and pupil response amplitude, we measured the normalized cross-correlation. In normal hearing mice, LA responses led pupil dilations during baseline recordings when sounds were novel, but the temporal correlation was significantly reduced as mice habituated to the sounds and testing protocol (**Figure 2e**). Following NIHL, the temporal correlation between LA activity and pupil-indexed arousal showed the opposite change: It was significantly increased rather than reduced (**Figure 2f**). Taken together, these findings demonstrate that acute peripheral sensory damage can disrupt the neural and autonomic habituation to familiar neutral sounds, converting the natural and adaptive suppression of response into neural sensitization and hyper-arousal.

### Noise-induced hearing loss disrupts discriminative threat learning

Affective processing is divided into two subdomains: arousal and valence^42^. The sustained elevation of LA responses and pupil dilations after NIHL changes in arousal regulation but did not address whether NIHL also disrupted the evaluation of sound valence. To examine this question, we employed discriminative threat conditioning (DTC), a Pavlovian learning protocol that presents a repeating sequence of relatively complex frequency modulated (FM) sounds that either always (CS+) or never (CS-) predict the delayed onset of an aversive primary reinforcer. Whereas auditory fear learning with simple sounds does not require neocortex, associative threat memories acquired through DTC depend upon higher order fields of the auditory cortex (HO-AC) and, even more specifically, their descending projections to the LA^26,32,43^.

After completing the 7-day passive listening assessment described above, NIHL and sham mice subsequently underwent a DTC protocol consisting of two Habituation sessions, three Conditioning sessions, a Recall session, and a final Extinction session (**Figure 3a**). During Habituation, Recall, and Extinction sessions, head-fixed mice were placed on a force transduction platform and presented with trains of upward or downward FM sweeps (constrained to the 5-11.3kHz range of intact hearing for all mice). Conditioning sessions occurred in a different sensory context, featuring interleaved trains of CS+ sweeps that co-terminated with a mild foot shock and CS-FM sweep trains that did not. The assignment of CS+ and CS- to upward vs downward FM sweeps was counterbalanced across mice (**Figure 3b**).

**Figure 3.**
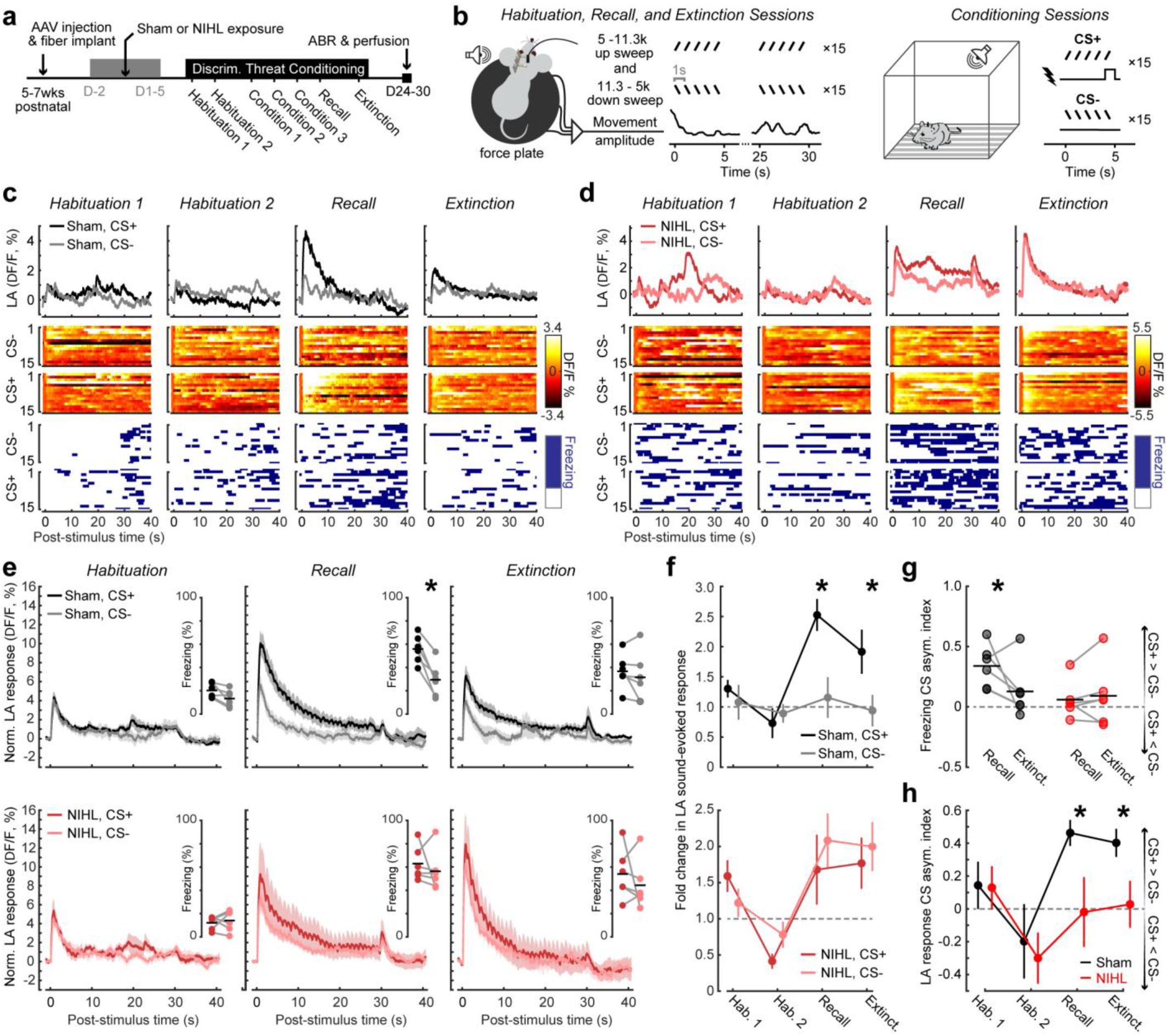
After NIHL, LA responses and freezing behavior generalizes across threatening and non-threatening sounds. (a) Experimental timeline. Discriminative threat conditioning (DTC) included 7 sessions of Habituation, Conditioning, Recall, and Extinction, beginning approximately one week following sham or NIHL in the same mice described in prior figures. (b) *Left*: Habituation, Recall, and Extinction sessions were performed in head-fixed mice resting atop a force plate that tracked body movements. Conditioned stimuli were 30s trains of upwards (5–11.3 kHz) and downwards (11.3–5 kHz) frequency modulated tone sweeps. *Right*: Conditioning sessions were performed in a different sensory context. Each session presented 30 interleaved 5s trials of one FM sweep direction that co-terminated with a mild foot shock (CS+) and the other sweep direction that did not (CS−). Assignment of upward vs downward sweeps to CS+/CS- was counterbalanced across mice. (**c–d**) LA responses and freezing behavior during Habituation, Recall, and Extinction sessions from a representative sham (c) and NIHL (*d*) mice. *Top*: mean LA GCaMP responses (ΔF/F, %) to CS+ and CS− across sessions. *Middle*: heatmaps of single-trial LA responses. *Bottom*: Freezing behavior quantified from the force plate signal for each corresponding trial. (e) Mean ± SEM LA GCaMP responses elicited by CS+ and CS- FM sweep trains for sham (*top,* N=6) and NIHL (*bottom*, N=6) groups. *Inset:* Freezing behavior elicited by the CS+ and CS- for each mouse (individual data points) and group means (horizontal black lines). Freezing was significantly greater for the CS+ than CS- in the Recall session for sham mice (paired t-test, T = 6.0, p = 0.002) but no significant differences were found for the remaining five comparisons (p > 0.2 for all). (f) Mean ± SEM fold change in LA evoked response amplitude to CS+ and CS- stimuli (relative to Habituation_mean_). LA responses were significantly greater for the CS+ than CS- after conditioning, whereas discriminative LA plasticity was not observed in NIHL mice (LME with stimulus type, session, and group as fixed factors; stimulus × session × group interaction, F = 2.87, p = 0.04). Asterisks indicate significant differences with post-hoc pairwise comparisons between CS+ and CS− for the Recall and Extinction tests in sham mice (p < 0.01 for both) but significant differences were not found in other conditions (p > 0.08 for all). (g) Freezing discriminability index (CS+ − CS−)/(CS+ + CS−) at Recall and Extinction. Sham mice showed more selective and extinguishing threat memory than NIHL mice (LME restricted to Recall and Extinction with session as a repeated measure and group as a between-subjects factor; main effect of group, F = 6.54, p = 0.018; main effect of session, F = 7.05, p = 0.015; session × group interaction, F = 4.66, p = 0.043). Pairwise comparisons (recall vs. extinction): sham, F = 7.05, p = 0.015; NIHL, F = 0.16, p = 0.69. Circles, individual mice; horizontal lines, group means. (h) Mean ± SEM LA discriminative plasticity index (CS+ − CS−)/(CS+ + CS−) at Recall and Extinction. Sham mice showed significantly greater discriminative plasticity after DTC than NIHL mice (LME restricted to recall and extinction sessions with session as a repeated measure and group as a between-subjects factor; main effect of group, F = 7.05, p = 0.014). Asterisks indicate significant between-group differences with post-hoc pairwise comparisons between sham and NIHL mice at the Recall (p=0.038) and Extinction (p=0.029).

In sham-exposed mice, LA GCaMP responses and freezing behavior were selectively enhanced for the CS+ stimulus during the Recall session but were subsequently reduced during Extinction (**Figure 3c and 3e, top**). NIHL mice showed a different pattern: LA responses and freezing behavior were indiscriminately enhanced for the CS+ and CS- during Recall and remained elevated in the Extinction session (**Figure 3d, and 3e, bottom**). At the group level, control LA responses to the CS+ were significantly enhanced during Recall and Extinction whereas CS- responses were unchanged (**Figure 3f, top**). In NIHL mice, LA responses to the CS+ and CS- were equivalently enhanced in Recall and Extinction sessions (**Figure 3f, bottom**). In sham mice, behavioral discrimination co-occurred with stimulus-selective LA responses: a CS+ vs CS- asymmetry index revealed significantly greater differential freezing to the CS+ stimulus during Recall than Extinction, whereas NIHL mice showed generalized freezing that failed to extinguish^44^ (**Figure 3g**). Applying the same CS+ vs CS- asymmetry index to LA responses confirmed that threat stimulus selectivity was significantly reduced compared to sham controls in the Recall and Extinction sessions (**Figure 3h**).

We next asked whether the impaired LA discrimination of CS+ and CS- stimuli after NIHL reflected degraded sensory discriminability. Using a stimulus-specific adaptation paradigm^45^, we found robust neural discriminability for upward and downward FM sweeps in NIHL mice (**Supplementary Figure 1**), indicating that the deficit more likely reflected overgeneralized threat learning than a failure to distinguish the low-level sensory features of the stimuli^44^.

### Acute activation of cortical PVNs suppresses sound-evoked LA responses

The LA receives convergent auditory input from thalamic and cortical sources. Because disinhibition and excess central gain are most prominently expressed in AC, we asked whether direct activation of AC PVNs was sufficient to suppress LA hyperresponsivity (**Figure 4a**)^10,35,46^. To test this, we expressed a somatically restricted red-shifted depolarizing opsin (Chrimson) or a control fluorophore (mCherry) in AC PVNs and investigated the acute effects of AC inhibitory activation on LA responses before and after NIHL (**Figure 4b**).

**Figure 4.**
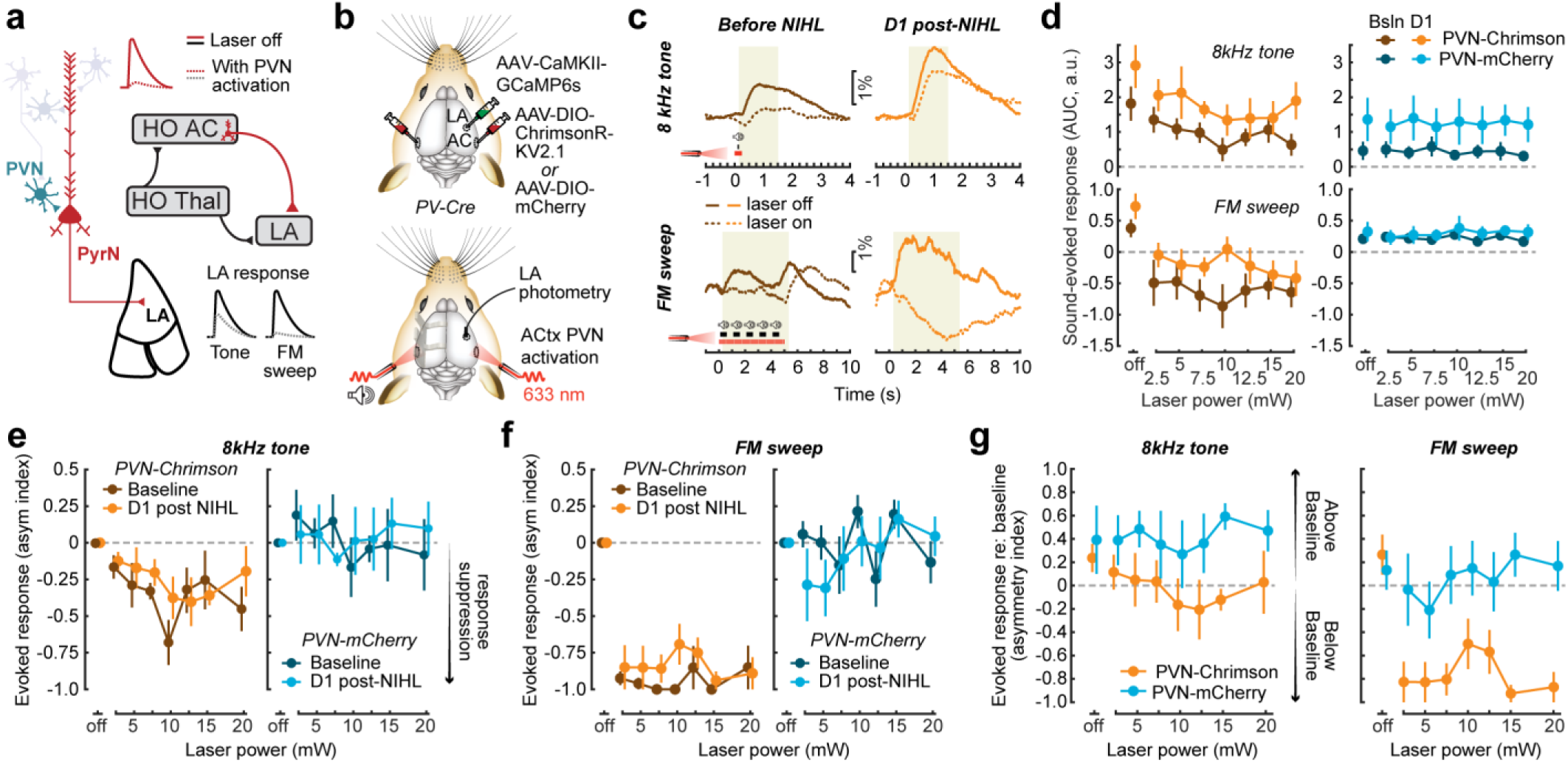
Attenuation of LA hyperresponsivity during activation of PVN inhibitory neurons in auditory cortex. (a) Circuit schematic illustrating the hypothesis that direct activation of parvalbumin-expressing inhibitory interneurons (PVNS) in the higher-order auditory cortex (HO-AC) would reduce LA hyperresponsivity after NIHL for simple (tones) and more complex (FM sweeps) sounds. (b) *Top:* PV-Cre received injections of AAV-CaMKII-GCaMP6s in the right LA, and injections of either a Cre-dependent somatically restricted red-shifted depolarizing opsin in the (AAV-DIO-ChrimsonR-KV2.1) or a Cre-dependent control fluorophore (or AAV-DIO-mCherry control) into the left and right auditory cortex (AC). *Bottom:* LA fiber photometry measurements were performed during the combined presentation of auditory stimuli and bilateral laser illumination. Recordings were performed in each mouse during two baseline sessions (D-2 and D-1) and one session occurring a day after NIHL (D1). (c) Example LA GCaMP responses to 60 dB SPL 8 kHz tones (*top*) and FM sweeps (*bottom*) in a representative PVN-Chrimson mouse with (dashed) and without (solid) AC PVN activation, before NIHL (*left*) and on D1 post-NIHL (*right*). Red bars indicate the timing of the brief continuous laser pulse and tone pip (*top*) or longer duration 20 Hz pulsed laser stimulation and FM sweep train (*bottom*). Yellow shading denotes the response analysis window. Scale bars, 1% ΔF/F. (d) Mean ± SEM sound-evoked LA responses to 60 dB SPL tones (*top*) and FM sweep trains (*bottom*) were computed from the area under the curve (AUC) of the GCaMP response during the response window shown in (*c*). Measurements were performed before (darker colors) and after (lighter colors) NIHL. Tone-evoked responses were reduced – and FM sweep responses eliminated - across a range of laser powers in PVN-Chrimson mice (N = 5; *left*) but not in PVN-mCherry (N = 6; *right*). Dashed horizontal gray bar = pre-stimulus activity level (i.e., no response). (e) Mean ± SEM suppression of sound-evoked responses in LA were quantified independently for each session with an asymmetry index referenced to the laser off condition (Power X – Off) / (Power X + Off). *Left*: PVN-Chrimson mice (N = 5; *left*) showed significantly greater suppression than mCherry controls (N = 6; *right*; LME with group and power as fixed effects and animal as random intercept; main effect of group, F = 9.12, p = 0.011). A separate model restricted to PVN-Chrimson mice confirmed power-dependent suppression (main effect of power, F = 3.38, p = 0.003) that did not differ across sessions (main effect of session, F = 0.81, p = 0.37). No effect of power or session was observed in mCherry controls (power, F = 0.34, p = 0.93; session, F = 0.10, p = 0.75). (f) As per *e,* but for FM sweep trains. PVN-Chrimson mice (*left)* showed significantly greater suppression than mCherry controls (*right;* LME with group and power as fixed effects and animal as random intercept; main effect of group, F = 209.88, p = 8.1 × 10⁻³²; main effect of power, F = 4.93, p = 4.2 × 10⁻⁵). A separate model restricted to PVN-Chrimson mice revealed significant effects of power (F = 32.58, p = 2.6 × 10⁻²⁰) and session (F = 5.77, p = 0.019). No effect of power or session was observed in mCherry controls (power, F = 1.38, p = 0.22; session, F = 1.34, p = 0.25). (g) Mean ± SEM sound-evoked response referenced to the pre-NIHL baseline laser off condition with an asymmetry index (Power X – Off_Baseline_) / (Power X + Off_Baseline_). Positive values indicate LA response sensitization after NIHL, negative values indicate LA responses are suppressed below baseline values, and a value of 0 indicates LA responses are restored to baseline values. Hyperresponsivity after NIHL was confirmed in the laser off condition for both tones (F = 5.05, p = 0.046; *left*) and FM sweeps (F = 7.94, p = 0.017; *right*), with both groups pooled (N = 11). Optogenetic activation of AC PVNs in PVN-Chrimson mice (N = 5) significantly suppressed LA responses to pre-NIHL levels or below for tones (F = 6.91, p = 0.013) and FM sweeps (F = 60.47, p = 3.9 × 10⁻⁹), while mCherry controls (N = 6) showed no suppression (tones, F = 0.058, p = 0.81; FM sweeps, F = 0.237, p = 0.63).

In the absence of cortical PVN activation, both low-frequency tones and FM sweeps evoked robust LA responses that were sensitized one day after NIHL (Figure 4c, solid traces). Activating AC PVNs during sound presentation suppressed responses to simple tone burst stimuli and eliminated LA responses to more complex FM sweep stimuli (**Figure 4c**). We titrated the degree of AC PVN activation by varying the laser power and observed a graded reduction in tone-evoked LA responses before and after NIHL (**Figure 4d top, Figure 4e**) and a power-independent elimination of FM sweep-evoked responses (**Figure 4d bottom, Figure 4f**). These effects were specific to PVN activation as bilateral laser activation had no effect on LA responses before or after NIHL in mCherry control mice (blue data points in **Figures 4d-f**).

To determine whether PVN activation restored LA responses toward the pre-NIHL state, we referenced each post-NIHL response to that mouse’s own baseline rather than to the laser-off condition. For this analysis, a positive asymmetry index value indicated LA sensitization after NIHL, negative values indicated that LA responses were suppressed below baseline, and a value of zero indicated that LA response were equivalent to measurements made before NIHL. We found that low levels of AC PVN activation restored tone-evoked LA responses to baseline values and completely suppressed FM-evoked LA responses, suggesting that AC PVN activation was sufficient to eliminate LA hyperresponsivity after noise-induced cochlear damage (**Figure 4g**).

### PVN gamma stimulation durably potentiates cortical inhibition and suppresses sound-evoked activity

PVN activation can, under some conditions, induce lasting changes in local circuit function that outlast the stimulation period^47,48^. For example, we recently showed that a single bout of PVN gamma frequency stimulation sustainably dampened sound-evoked population activity and enhanced PVN-mediated feedforward inhibition in the primary auditory cortex, and behaviorally desensitized mice to loudness for one week^18^. Whether these effects extend to the HO-AC and beyond to brain regions outside the central auditory pathway is unknown. If 40-Hz PVN stimulation could durably potentiate PVN feedforward inhibition and dampen sound-evoked responses in HO-AC neurons, it could represent an approach for normalizing maladaptive LA plasticity after NIHL (**Figure 5a**).

**Figure 5.**
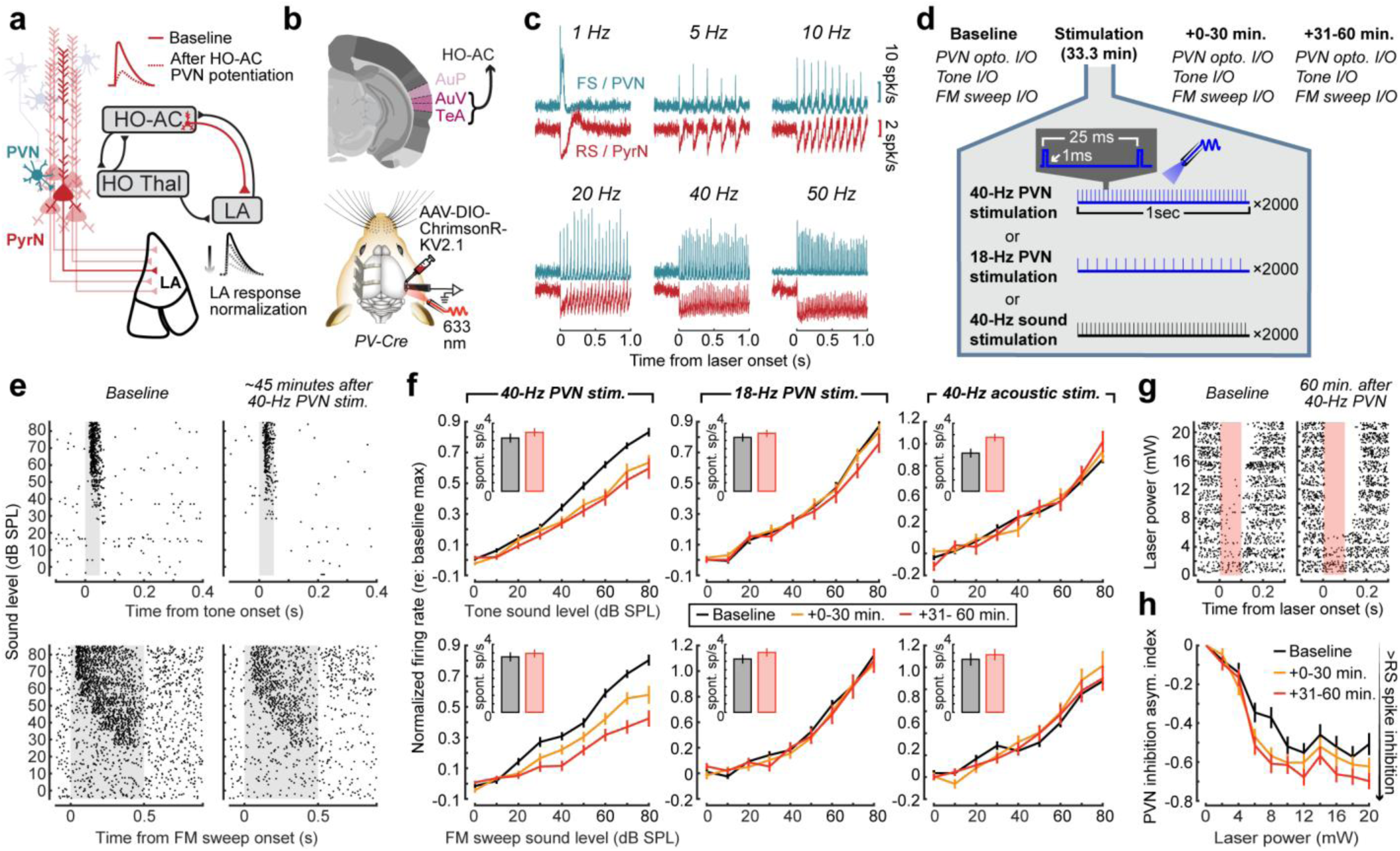
A sustained period of 40-Hz PVN activation durably suppresses sound-evoked spiking and potentiates PVN-mediated inhibition in higher-order auditory cortex. (a) Circuit schematic illustrating the hypothesis that potentiating parvalbumin-expressing inhibitory neurons (PVNs) in higher-order auditory cortex (HO-AC) could durably reduce LA hyperactivity. PyrN = pyramidal neuron. (b) *Top*: Coronal section depicts three regions of the auditory cortex, of which AuV and and TeA are operationally defined here as the higher-order auditory cortex (HO-AC). Nomenclature and reference image are adapted from the Allen Institute for Brain Science. *Bottom:* AAV-DIO-ChrimsonR-KV2.1 was expressed in auditory cortex PVNs in PV-Cre mice. Optogenetic stimulation and extracellular single unit recordings were performed in the right HO-AC of unanesthetized head-fixed mice. (c) Mean fast spiking (FS) / PVN (n = 16, teal) and regular spiking (RS) / PyrN (n = 260, red) unit PSTHs in HO-AC during 1 s optogenetic PVN stimulation at 1–50 Hz. Scale bars, 10 spk/s (FS) and 2 spk/s (RS). (d) Experimental design for 40-Hz PVN stimulation, 18-Hz PVN stimulation, or 40-Hz acoustic stimulation. Each protocol delivered continuous stimulation for 2000 seconds (33.3 min). RS spiking input-output functions were measured in response to laser pulses of varying power, tone bursts of varying sound levels, and FM sweeps of varying sound levels at three time points: Baseline, 0–30 min following the cessation of the stimulation period, and 31–60 min after the stimulation period ended. (e) Raster plots from an example RS unit showing reduced tone-evoked (*top*) and FM sweep-evoked (*bottom*) spiking across sound levels approximately 45min after the 40-Hz PVN stimulation period ended (*right*). (f) Mean ± SEM normalized RS spike rate sound level growth functions for tones (*top*) and FM sweeps (*bottom*) were referenced to maximum response at baseline. *Inset:* Spontaneous RS spike rates measured at baseline and 31-60min. *Left column:* After 40-Hz PVN stimulation, tone- and FM sweep-evoked responses were significantly suppressed (N = 4 mice, n = 137 tone/109 FM sweep units; LME with time as a repeated measure; evoked tone, F = 15.23, p = 4.18 × 10⁻⁷; evoked FM sweep, F = 49.83, p = 1.36 × 10⁻¹⁹). Pairwise comparisons confirmed suppression at >60 min for both tones (F = 18.85, p = 2.0 × 10⁻⁵) and FM sweeps (F = 83.00, p = 5.7 × 10⁻¹⁷), with no change in spontaneous rates (tone, p = 0.17; FM sweep, p = 0.47). *Middle column:* No significant suppression was observed after 18-Hz PVN stimulation (N = 2, n = 108/80 units; tone, p = 0.061; FM sweep, p = 0.41). *Right column:* After 40-Hz acoustic stimulation, no significant evoked response changes were observed (N = 2, n = 102/102 units; tone, p = 0.77; FM sweep, p = 0.78), though spontaneous rates increased for tones (F = 17.71, p = 3.9 × 10⁻⁵). (g) Example RS unit rasters showing PVN-mediated inhibition of spontaneous spiking across laser powers at baseline (*left*) and approximately 60 min after 40-Hz PVN stimulation (*right*). Red shading indicates the laser-on period. (h) Mean ± SEM PVN inhibition asymmetry index as a function of laser power computed as (FR_XmW − FR_0mW) / (FR_XmW + FR_0mW), where negative values indicate increasing PVN-mediated inhibition (N = 4 mice, n = 110 RS units). PVN-mediated inhibition was significantly enhanced after 40-Hz PVN stimulation (LME with time and power as fixed effects and animal as random intercept; time × power interaction, F = 3.72, p = 0.025). Pairwise comparison confirmed stronger inhibition at >60 min relative to baseline (F = 32.24, p = 1.3 × 10⁻¹⁴).

To test this, we made acute extracellular recordings of regular-spiking (RS/pyramidal) and fast-spiking (FS/PVN) units from HO-AC in unanesthetized head-fixed mice with high-density silicon probes (**Figure 5b**). Optogenetic activation of Chrimson-expressing PVNs entrained RS and FS spiking across a wide range of pulse rates, with FS units faithfully following each pulse and RS units showing time-locked inhibition-rebound patterns (**Figure 5c**). We then compared three stimulation protocols, each delivering sustained activation over a 33.3 minute period: 40-Hz PVN optogenetic stimulation, 18-Hz PVN optogenetic stimulation, and 40-Hz acoustic stimulation. Sound-evoked responses (tones and FM sweeps), PVN-mediated inhibition, and spontaneous spiking were measured at baseline, +0–30 min, and 31-60 min following the end of stimulation (**Figure 5d-e**).

Gamma stimulation of HO-AC PVNs durably suppressed tone- and FM sweep-evoked RS unit spiking across a wide range of sound levels without significantly affecting the spontaneous spiking rate (**Figure 5f, left**). This short-term plasticity was only observed with PVN gamma stimulation, as neither 18-Hz PVN stimulation nor 40-Hz acoustic stimulation had a significant effect on sound-evoked activity (**Figure 5f, middle and right**). While direct measures of synaptic inhibition are not possible with extracellular recordings, we indirectly measured PVN-mediated feedforward inhibition by quantifying the suppression of RS unit spontaneous spiking during PVNs activation with laser pulses of increasing power (**Figure 5g**). We found that PVN-mediated inhibition was significantly enhanced after 40-Hz PVN stimulation, with a significant time × power interaction confirming that inhibition strengthened progressively with laser power, especially at longer wait times following the period of gamma stimulation (**Figure 5h**). Taken together, these recordings show that 40-Hz stimulation of HO-AC PVNs durably strengthened feedforward inhibition and suppressed sound-evoked spiking in putative pyramidal neurons, supporting its potential utility as an approach to reverse disruptions in affective sound processing following NIHL.

### Potentiating cortical inhibition eliminates LA hyperresponsivity

We next addressed whether enhancing cortical inhibition via a bout of 40-Hz PVN stimulation could durably reverse LA hyperresponsivity over several days in a manner analogous to the acute effects cortical PVN activation (Figure 4). These experiments featured three distinct phases of testing for each mouse: 1) Two days of baseline LA recordings before NIHL (D-2 and D-1); 2) One recording session after NIHL (D2); 3) Three days of recordings following a single 33min bout of bilateral HO-AC PVN 40-Hz stimulation (D3-D5; **Figure 6a**). As illustrated in an example mouse, the initial sensitization of LA sound level growth functions following NIHL was subsequently reversed after bilateral PVN 40-Hz stimulation to even lower levels than observed as baseline, on par with the habituation observed in mice with normal hearing (**Figure 6b and 6c, right**). By contrast, a control NIHL mouse expressing mCherry in cortical PVNs exhibited stable LA hyperresponsivity after 40-Hz stimulation (**Figure 6b and 6c, left**).

**Figure 6.**
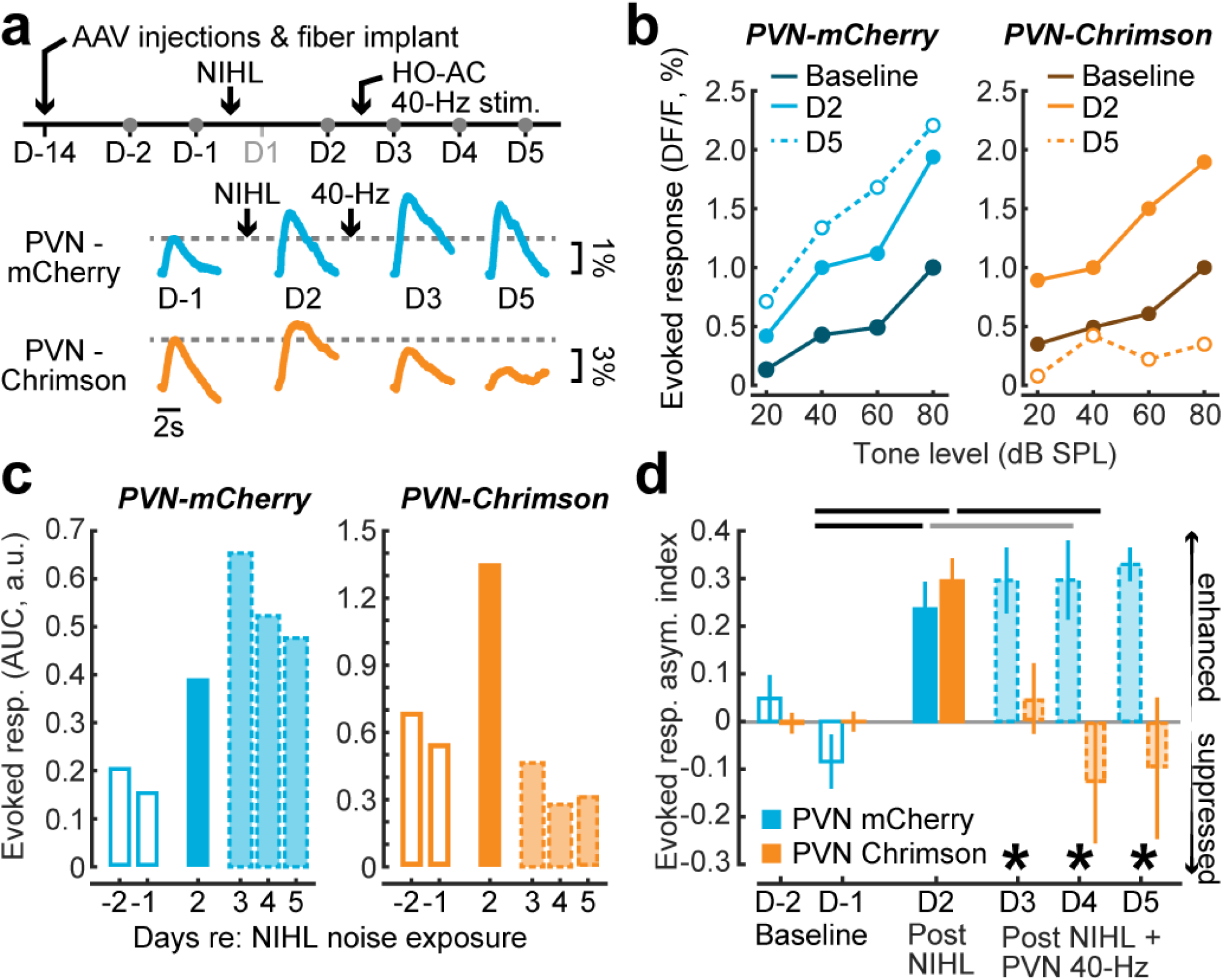
Cortical inhibitory potentiation sustainably reverses LA hyperresponsivity after noise-induced hearing loss. (a) *Top:* Experimental timeline. LA GCaMP recordings were compared across two baseline sessions (D-2, D-1) and four sessions after NIHL (D2-D5). Bilateral 40-Hz PVN stimulation was performed immediately after the conclusion of the D2 recording session. Thus, LA responses could be measured at three distinct phases: Before NIHL (D-2, D-1), after NIHL (D2), and after NIHL + HO-AC PVN gamma stimulation (D3-5). *Below*: Tone-evoked LA GCaMP responses (8 kHz at 80 dB SPL) from an example PVN-mCherry (*top*) and PVN-Chrimson (*bottom*) mouse across sessions illustrates the the response normalization after 40-Hz PVN stimulation but the stable sensitization in mCherry mice. Scale bars, 1% ΔF/F (mCherry) and 3% ΔF/F (Chrimson); time bar, 2 s. (b) Tone-evoked LA sound level growth functions (peak – baseline, ΔF/F, %) at baseline (dark blue/brown), D2 post-NIHL (cyan/orange), and D5 (dashed) for an example PVN-mCherry (*left*) and PVN-Chrimson (*right*) mouse. (c) Evoked responses from the same example mice were collapsed to a single value by calculating the area under the curve (AUC) for 40-80 dB SPL sound levels. AUC values are provided for all six photometry sessions. (d) Mean ± SEM Evoked response asymmetry index was quantified as ([Session X – Baseline_mean_] / [Session X + Baseline_mean_]), where negative values indicate suppression relative to baseline and positive values indicate hyper-responsiveness (sensitization). LA responses were stably sensitized after NIHL in PVN-mCherry mice (N=6) but sensitization was reversed after PVN gamma stimulation in PVN-Chrimson mice (N=5; LME with session as a repeated measure and group as a between-subjects factor; main effect of session, F = 10.69, p = 3.6 × 10⁻⁷; session × group interaction, F = 8.98, p = 2.9 × 10⁻⁶). Post-hoc pairwise comparisons with Holm–Bonferroni correction revealed that both groups showed NIHL-induced hyperactivity on D2 ([D-2, D-1] vs. D2: mCherry, p = 3.2 × 10⁻⁴; Chrimson, p = 6.7 × 10⁻⁵). Following 40-Hz PVN stimulation, LA responses were suppressed back toward baseline in PVN-Chrimson mice (D2 vs. [D3, D4, D5], p = 6.38 × 10⁻⁶, black horizontal lines) but not in mCherry controls (p = 0.23, gray horizontal line). Between-group comparisons confirmed divergence beginning on D3 (D3, p = 0.03; D4, p = 9.0 × 10⁻⁵; D5, p = 1.88 × 10⁻⁴, asterisks), but no significant group differences beforehand (p = 1.0 for each).

We quantified the change in LA responsivity using an evoked response asymmetry index referenced to each mouse’s pre-NIHL baseline where positive and negative values indicate hyperresponsivity and suppression, respectively (**Figure 6d**). PVN-mCherry and PVN-Chrimson groups both showed significant NIHL-induced hyperactivity on D2, replicating the effect of NIHL on LA response sensitization in a separate cohort of PV-Cre mice with bilateral virus expression. Following 40-Hz stimulation, LA responses were suppressed back toward baseline in PVN-Chrimson mice but not in mCherry controls. Between-group comparisons confirmed that the groups diverged beginning on D3, with the Chrimson group showing progressively greater suppression through D5.

### Cortical inhibitory potentiation reverses auditory hyper-arousal

Our initial characterization study found that effects of NIHL extended beyond the sensory phenotype of neural hyperresponsivity to include changes in affective arousal and valence processing studied by sound-evoked pupil dilations (Figure 2) and DTC (Figure 3), respectively. Having demonstrated that LA hyperresponsivity can be sustainably reversed for at least three days after PVN 40-Hz stimulation, we next asked whether affective arousal and valence processing were also changed to more closely resemble the pre-NIHL state. Sound-evoked pupil responses were comparable between PVN-mCherry and PVN-Chrimson groups at baseline, confirming equivalent autonomic reactivity before any manipulation (**Figure 7a,b, left**). On D2 after NIHL, both groups demonstrated sensitized pupil-indexed arousal sound level growth functions, replicating the hyperarousal phenotype observed in the initial cohort of wildtype mice in a second cohort of PV-Cre mice with bilateral virus injections (**Figure 7a,b, middle**). From here the groups diverged: Pupil-indexed arousal reversed to baseline levels on D3-5 after 40-Hz PVN stimulation but exhibited sustained hyper-arousal in mCherry controls (**Figure 7a,b, right**).

**Figure 7.**
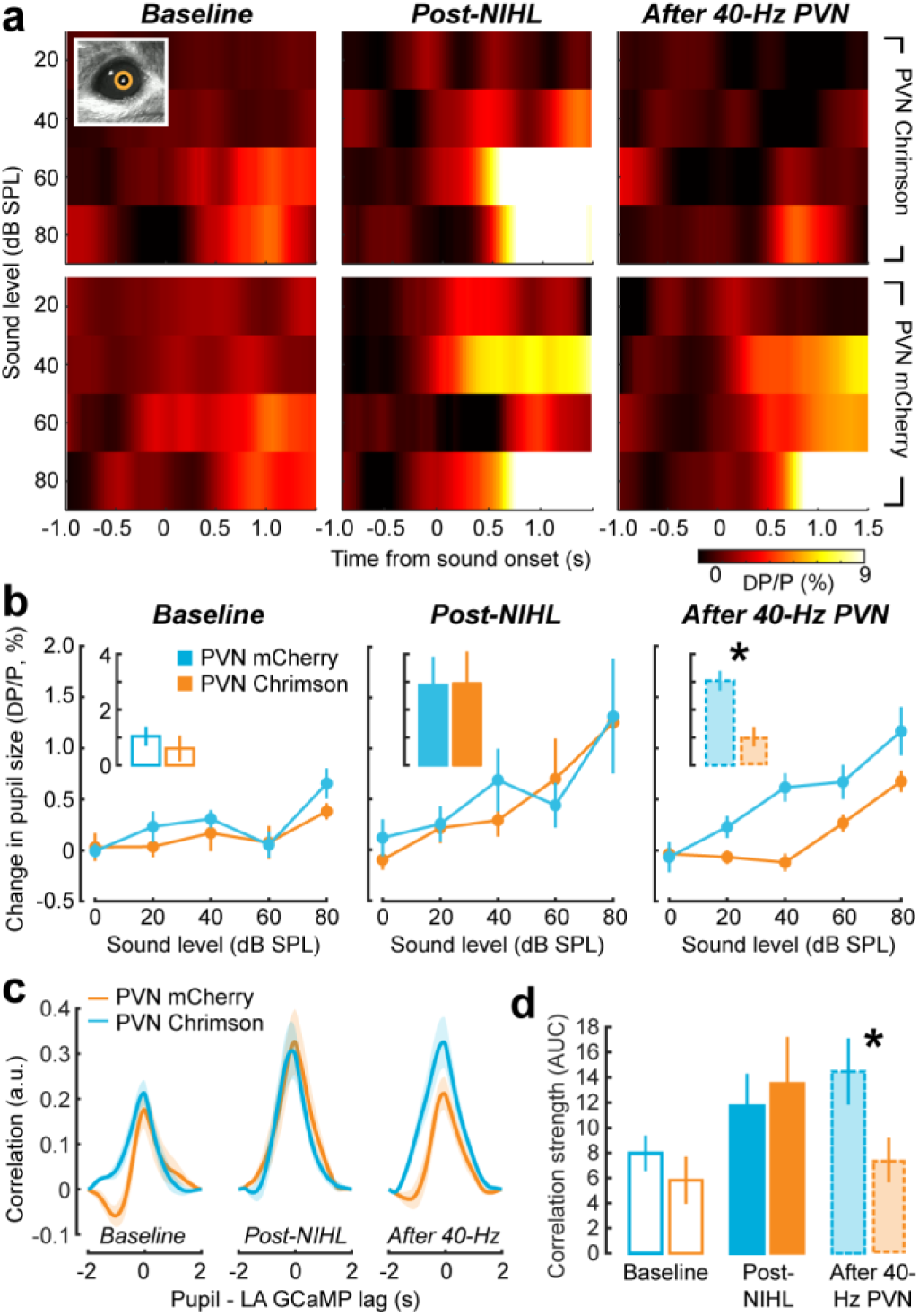
Cortical inhibitory potentiation reverses pupil-indexed hyperarousal and abnormal pupil–LA coupling. (a) Pupillometry was performed concurrently with LA fiber photometry recordings. Heatmaps depict mean sound-evoked pupil dilation by 8kHz tones (ΔP/P, %) as a function of sound level for a representative PVN-Chrimson (*top)* and PVN-mCherry (*bottom*) mouse at baseline (*left*), D2 post-NIHL (*middle*), and after 40-Hz PVN stimulation (*right*). *Inset*: example pupil video frame with tracking overlay. (b) Mean ± SEM sound-evoked change in pupil size as a function of sound level for PVN-mCherry (blue, N = 6) and PVN-Chrimson (orange, N = 5) mice at baseline (*left*), post-NIHL (*middle*), and after 40-Hz PVN stimulation (*right*). *Insets*: mean pupil response averaged across 40–80 dB SPL. Pupil responses were compared using an LME with session as a repeated measure and group as a between-subjects factor (main effect of session, F = 10.12, p = 1.8 × 10⁻⁴; session × group interaction, F = 4.57, p = 0.014). Pairwise comparisons revealed no group difference at baseline (p = 0.81) or post-NIHL (p = 0.93), but a significant divergence after 40-Hz PVN stimulation (p = 4.8 × 10⁻⁵, asterisk). (c) Mean ± SEM normalized cross-correlation between LA GCaMP and pupil signals measured in PVN-mCherry (N=6) and PVN-Chrimson (N=5) mice at baseline (*left*), D2 post-NIHL (*middle*), and D3-D5 40-Hz stimulation (*right*). (d) Mean ± SEM LA-pupil temporal coupling, quantified as the AUC of positive cross-correlation values in PVN-mCherry (N=6) and PVN-Chrimson (N=5) groups. LME with session as a repeated measure and group as a between-subjects factor; main effect of session, F = 3.90, p = 0.026). Pairwise comparisons revealed no group difference at baseline (p = 0.31) or post-NIHL (p = 0.60), but a significant divergence after 40-Hz PVN stimulation (p = 0.003, asterisk).

We then examined whether 40-Hz PVN stimulation also reversed the abnormally strong temporal coupling between LA activity and pupil dilation observed after NIHL (see Figure 2e-g). Normalized cross-correlations between the pupil and LA GCaMP signals showed similar coupling profiles for both groups at baseline and post-NIHL, replicating the NIHL-induced enhancement of LA–pupil coupling (**Figure 7c and 7d, left and middle**). After 40-Hz PVN stimulation, coupling strength reverted to baseline levels while mCherry controls maintained elevated coupling (**Figure 7c and 7d, right**). Thus, 40-Hz PVN stimulation normalized both sound-evoked pupil dilation and LA–pupil coupling after NIHL.

### Cortical inhibitory potentiation reinstates selective behavioral threat learning and recall

We next asked whether these corrections translate into recovery of the behavioral and neural phenotype of abnormal auditory valence processing after NIHL: overgeneralized threat memory recall and impaired extinction (see Figure 3). These experiments used the same 7-day Habituation, Conditioning, Recall, and Extinction protocol used in the wildtype cohort, with the exception of a second 33min bout of 40-Hz stimulation occurring 1-2 hours before the first Conditioning session (**Figure 8a**). Because 40-Hz stimulation preceded Conditioning, this design tests whether PVN gamma stimulation can prevent, rather than reverse, the maladaptive associative learning phenotype observed after NIHL.

**Figure 8.**
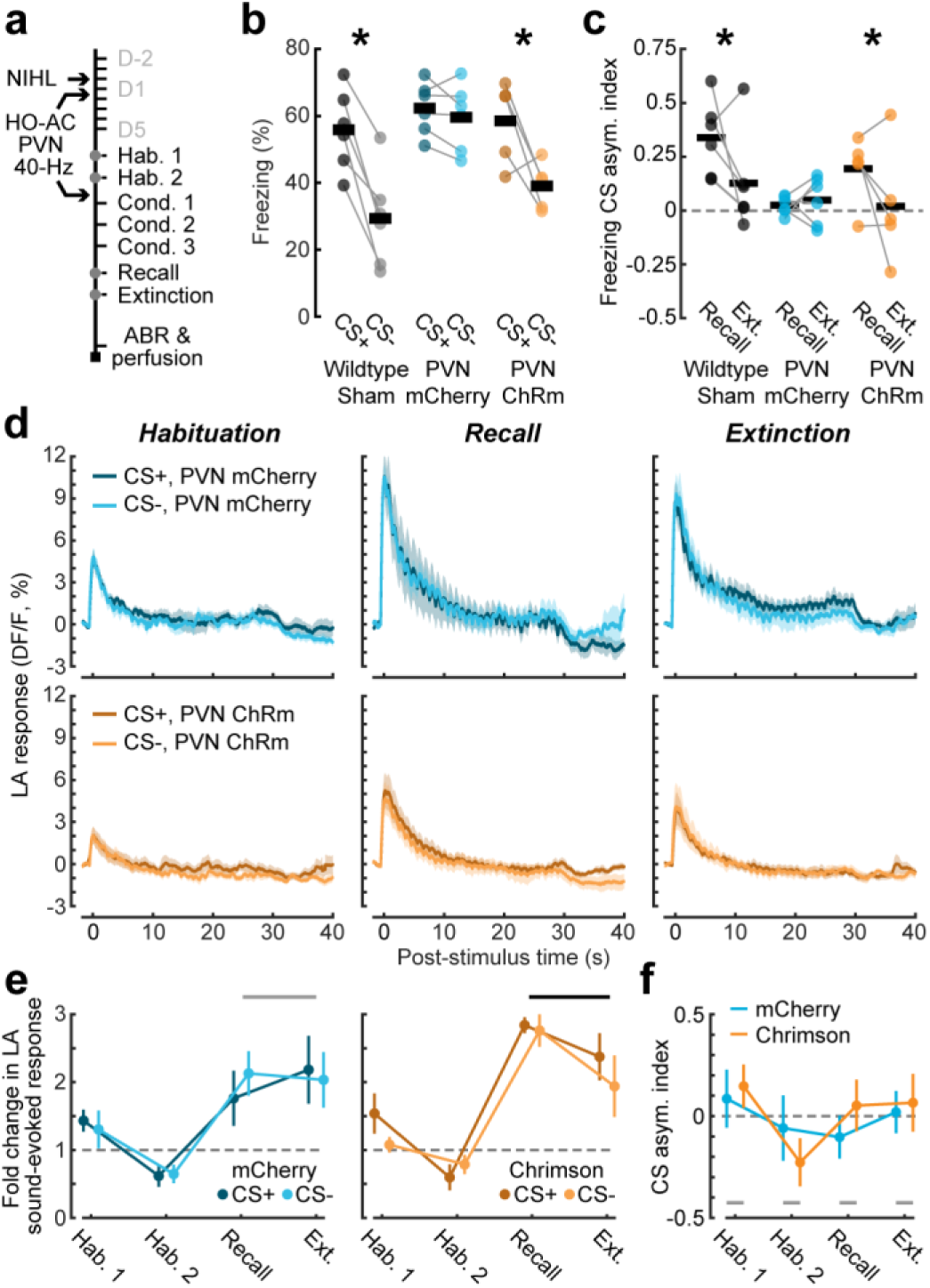
Cortical inhibitory potentiation reinstates discriminative threat learning but indiscriminate LA plasticity persists. (a) Experimental timeline. Discriminative threat conditioning (DTC) included 7 sessions of Habituation, Conditioning, Recall, and Extinction, beginning approximately one week following sham or NIHL in the same PV-Cre mice described in prior figures. The second bout of bilateral 33.3 minute 40-Hz PVN stimulation occurred 1–2 h before the first Conditioning session. (b) Freezing behavior elicited by the CS+ and CS- during the Recall session for each mouse (individual data points) and group means (horizontal black lines). Freezing in sham exposed wild type mice is replotted from Figure 3 for comparison to NIHL PVN-mCherry (N=6) and NIHL PVN-Chrimson (N=5) mice. Freezing to the CS+ stimulus was significantly greater than the CS- in sham-exposed and PVN-Chrimson mice (paired t tests; sham, t = 6.0, p = 0.002; Chrimson, t = 2.78, p = 0.049), but not in PVN-mCherry mice (t = 1.27, p = 0.26). (c) Freezing discriminability index (CS+ − CS−)/(CS+ + CS−) at Recall and Extinction. Sham and NIHL PVN-Chrimson mice showed more selective and extinguishing threat memory than NIHL PVN-mCherry mice (LME restricted to Recall and Extinction in PV-Cre mice with session as a repeated measure and group as a between-subjects factor; main effect of group, F = 6.54, p = 0.018; main effect of session, F = 7.05, p = 0.015; session × group interaction, F = 4.66, p = 0.043). Asterisks indicate significant pairwise differences (Recall vs. Extinction): Sham, p = 0.015; PVN-Chrimson (p = 0.039), PVN-mCherry (p = 0.75). (d) Mean ± SEM LA GCaMP responses elicited by CS+ and CS- FM sweep trains for PVN-mCherry (*top,* N=6) and PVN-Chrimson (*bottom*, N=5) groups. CS+ and CS- stimuli were 30s trains of upwards (5–11.3 kHz) and downwards (11.3–5 kHz) frequency modulated tone sweeps. (e) Mean ± SEM fold change in LA evoked response amplitude to CS+ and CS- stimuli (relative to Habituation_mean_) for PVN-mCherry (N = 6; *left*) and PVN-Chrimson (N = 5; *right*) mice. LME with stimulus and session as repeated measures and group as a between-subjects factor; main effect of session, F = 13.09, p = 3.8 × 10⁻⁷; stimulus, F = 1.43, p = 0.24; group, F = 0.08, p = 0.77; session × group × stimulus interaction, F = 0.27, p = 0.85). PVN-Chrimson mice showed a significant reduction in overall LA response magnitude between the Recall and Extinction sessions (LME with session as a repeated measure; F = 7.34, 15, p = 0.016, black horizontal line), but PVN-mCherry control did not (F = 0.32, p = 0.579; gray horizontal line). (f) Mean ± SEM LA discriminative plasticity index (CS+ − CS−)/(CS+ + CS−). Discriminative CS+/CS- plasticity was not observed across groups and test sessions (LME with session as a repeated measure and group as a between-subjects factor; main effect of session, F = 0.008, p = 0.92; session × group interaction, F = 0.70, p = 0.41). Gray bars indicate that pairwise differences between PVN-Chrimson and PVN-mCherry groups were not significant in any test session (p>0.32 for all).

At Recall, NIHL mice receiving 40-Hz PVN stimulation showed significantly greater freezing to the CS+ than CS− stimuli, on par with the selective threat memory recall observed in mice with normal hearing (**Figure 8b, left and right**). By contrast, NIHL mice with bilateral mCherry expression in PVNs exhibited non-selective freezing to CS+ and CS−, replicating the over-generalized threat memory recall observed in wildtype NIHL mice (**Figure 8b, middle**). The freezing CS asymmetry index confirmed this pattern: After PVN 40-Hz stimulation, NIHL mice demonstrated a selective threat memory at Recall that was significantly reduced during the Extinction session (**Figure 8c, left and right**). Mice expressing mCherry in PVNs exhibited similar levels of non-selective freezing in the Recall and Extinction sessions (**Figure 8c, middle**).

Unexpectedly, behavioral rescue was not accompanied by restoration of stimulus-selective bulk LA responses. Despite the recovery of behavioral discrimination and extinction, concurrently recorded LA responses remained similarly enhanced for threat-predicting and non-threat-predicting sounds in both groups. PVN-Chrimson and PVN-mCherry groups both exhibited non-discriminative enhancement of CS+ and CS− in the Recall session compared to Habituation (**Figure 8d and 8e**). The CS asymmetry index did not differ between groups or from zero (**Figure 8f**). The only significant difference observed between PVN-Chrimson and PVN-mCherry mice was a

reduction in overall LA response magnitude between the Recall and Extinction sessions. PVN 40-Hz stimulation therefore restored behavioral discrimination and extinction without significantly reinstating stimulus-selective LA responses.

## Discussion

Taken together, these experiments delineate a pathway from permanent noise-induced cochlear injury to affective dysfunction and demonstrate recovery through an intervention targeting cortical inhibitory networks.

Our prior work showed that NIHL weakened PVN-mediated inhibition drove hyperresponsivity in cortical excitatory neurons and loudness hypersensitivity⁵. Here we show that this cortical hyperactivity propagates to subcortical limbic and autonomic circuits: Following NIHL, spared low-frequency sounds elicited sustained LA hyperresponsivity, amplified sound-evoked pupil dilation, and strengthened the temporal coupling between LA activity and autonomic arousal (Figures 1–2). NIHL also disrupted discriminative threat learning — producing overgeneralized freezing for the CS+ and CS- sounds, reduced extinction, and indiscriminately enhanced LA responses to the CS+ and CS- (Figure 3). Acute optogenetic activation of auditory cortex PVNs suppressed LA hyperresponsivity, establishing a role for cortical inhibition in the regulation of LA sound responses (Figure 4).

Bouts of AC 40-Hz PVN stimulation sustainably potentiated intracortical inhibition and dampened sound-evoked cortical spiking in HO-AC units (Figure 5), eliminated LA hyperresponsivity (Figure 6), normalized pupil hyperarousal and LA–pupil coupling (Figure 7), and rescued both CS+/CS− behavioral discrimination and the capacity for threat memory extinction (Figure 8). These findings build on prior findings that focused on the sensory dimensions of hyperacusis by showing that cortical PVNs circuits also regulate the affective dimensions of hyperacusis, offering a control node to normalize aberrant downstream limbic and autonomic processing despite irreversible cochlear sensorineural damage.

A notable finding was that selective freezing and extinction were restored by 40-Hz PVN stimulation even though LA population responses were indiscriminately amplified to the CS+ and CS- sounds (Figure 8). Three features of LA circuit organization could explain this discrepancy. First, behavioral discrimination and extinction do not require reinstatement of LA selectivity per se — they instead require appropriate gating of amygdala output by downstream prefrontal and central amygdala circuits that regulate fear expression independently of LA response magnitude^49–54^; extinction in particular involves a shift in the balance of activity among defined amygdala neuronal populations rather than a silencing of LA responses^55^, and is therefore sensitive to the cortical and circuit context in which LA activity is embedded. When 40-Hz PVN stimulation enhanced cortical inhibitory tone, it may have restored these downstream gating mechanisms, allowing downstream circuits to extract differential stimulus value even without overt LA population selectivity. Consistent with this interpretation, PVN-Chrimson mice showed a significant reduction in overall LA response magnitude from Recall to Extinction — an effect absent in mCherry controls — suggesting that 40-Hz PVN stimulation re-engaged cortical feedback suppression of LA activity during extinction, even in the absence of restored stimulus selectivity. Second, LA population activity broadly tracks associative salience and arousal state rather than encoding stimulus identity with the precision required to determine behavioral output^22,56,57^; the large, non-discriminative responses seen after NIHL are consistent with elevated and non-specific gain within LA circuits, in which all salient sounds acquire equivalent associative weight. Third, CS selectivity in the LA under normal conditions is sparse and ensemble-based: fear conditioning recruits a minority of LA principal neurons whose synapses undergo associative plasticity, while simultaneously depressing responses in others — such that associative encoding is embedded at the ensemble level and is not reliably detectable in bulk population signals^55,58,59^; notably, disrupting this engram sparsity specifically within the LA is sufficient to convert specific threat memories into generalized ones^56,60^. The non-selective population responses observed here may therefore reflect a pathological reduction in engram sparsity that persisted despite behavioral rescue — or alternatively, normal engram structure may have been restored by 40-Hz PVN stimulation but remained below the resolution of bulk fiber recordings. These findings suggest that normalization of cortical gain, rather than reinstatement of LA response selectivity, may be sufficient to restore the dynamic range within which behavioral discrimination can emerge.

There are inherent limitations in our experimental approach that should be considered when evaluating these findings. First, although the convergence of cortical dampening (Figure 5) and LA normalization (Figure 6) in the same animals is consistent with a reduction in corticoamygdalar drive, we did not directly record from identified corticoamygdalar projection neurons or measure synaptic transmission along this pathway; whether corticoamygdalar neurons directly drive changes in LA responses, rather than arising through a parallel route, requires direct projection-specific interrogation. Additionally, because NIHL may also alter thalamic relay neuron activity, and the direct thalamo-amygdalar pathway contributes to LA tone-evoked responses, thalamic contributions to NIHL-induced LA hyperactivity cannot be excluded. The near-complete cortical dependence of FM-evoked LA responses suggests that the cortical route is the dominant pathway for the complex stimuli used in threat conditioning, but thalamic changes may contribute to the tone-evoked hyperactivity documented in Figures 1 and 6. This difference reinforces the greater cortical dependence of FM-evoked LA activity and suggests that reversing NIHL-induced LA hyperactivity to complex, cortically dependent stimuli may require less cortical intervention than reversing hyperactivity to stimuli with parallel subcortical access. Second, we cannot determine from the current data whether normalization of pupil hyperarousal following 40-Hz PVN stimulation was mediated through the LA or other parallel pathways through which the cortex can modulate brainstem regulators of the autonomic nervous system^61,62^. Third, the 40-Hz PVN stimulation was delivered prior to fear conditioning, making this a preventive rather than therapeutic intervention; whether the same protocol can reverse already-consolidated generalized fear memory remains an important open question, as the present design tested prevention rather than reversal of maladaptive associative learning. Importantly, the convergence of cortical, amygdala, autonomic, and behavioral normalization within the same animals argues against a trivial or regionally isolated explanation and supports a model in which cortical inhibitory tone regulates LA gain and affective sound processing as part of a coordinated systems-level response to cochlear injury. The precise synaptic and circuit mechanisms through which this occurs will require projection-specific recording and manipulation approaches to resolve fully.

More broadly, these findings offer a new perspective on the relationship between sensory injury and affective dysfunction. The high prevalence of anxiety, fear, and avoidance behaviors among patients with cochlear damage has long been recognized^33,63,64^, and autonomic measures such as sound-evoked pupil dilation independently predict psychiatric symptom severity^33^ — yet the neural substrate linking peripheral hearing loss to these affective sequelae has remained unknown. The present findings suggest that disinhibition of corticoamygdalar projections following cochlear injury may be a key intermediate step: elevated cortical gain, transmitted via direct auditory cortex projections to the LA, may alter limbic responsivity and autonomic reactivity to sound in ways that bias associative learning toward generalization and impair extinction. This account complements prevailing cortico-striato-thalamic gating models of tinnitus and hyperacusis^7–9^ by highlighting a direct corticoamygdalar pathway that may transmit cortical hyperactivity to limbic circuits, and extends the framework established by our previous study^18^ by suggesting that the perceptual and affective consequences of cochlear injury are linked by a shared alteration in cortical inhibitory tone — one that is amenable to correction through a single intervention. Notably, gamma-frequency entrainment of PV interneurons has now been shown to produce sustained beneficial effects across multiple domains of cortical dysfunction, from Alzheimer’s-related network disruption^48^ to post-stroke synaptic plasticity^47^, raising the possibility that patterned activation of cortical inhibitory circuits could represent a broader strategy for reversing maladaptive plasticity. Whether analogous corticoamygdalar mechanisms contribute to the affective dimensions of other sensory disorders — including chronic pain, visual loss, or age-related auditory decline — remains an important open question, but the present results suggest that the sensory cortex may be a surprisingly accessible control node for normalizing limbic dysregulation caused by peripheral injury.

## SUPPLEMENTAL FIGURES AND FIGURE LEGENDS

**Supplemental Figure 1.**
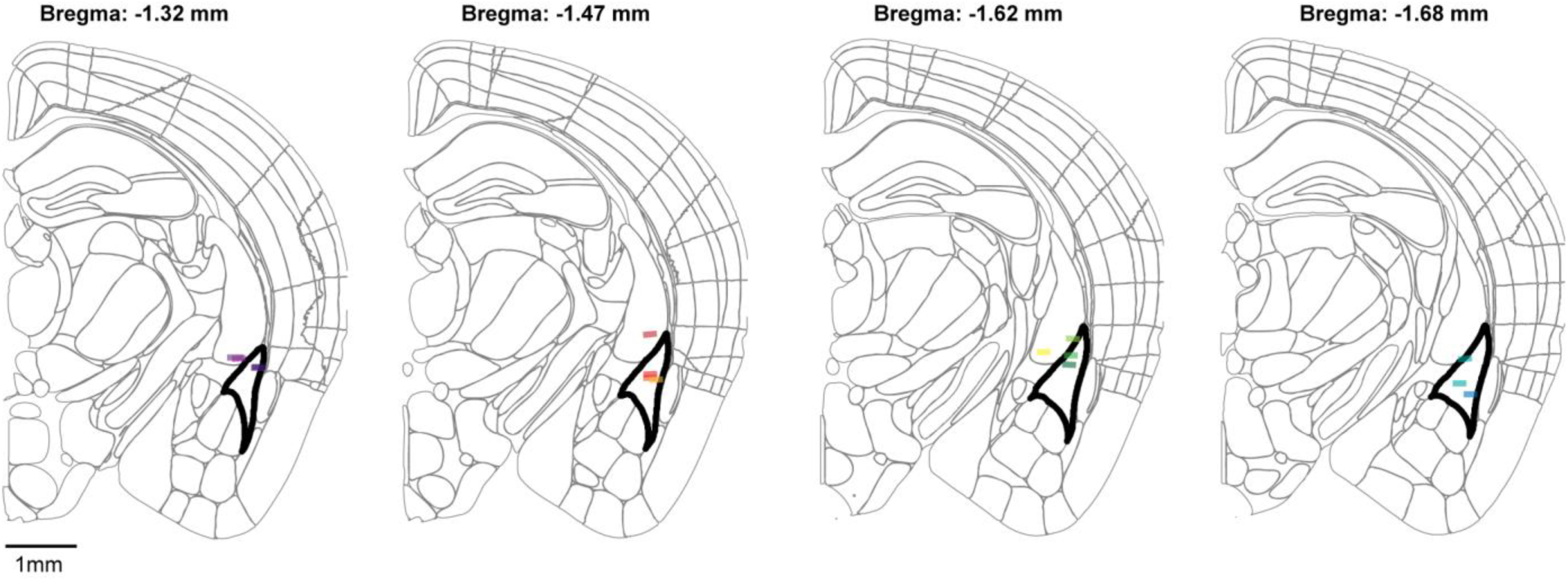
Histological verification of fiber tip locations. The center of each fiber tip location was identified in post-mortem coronal sections in wild type mice (N = 15, each identified by a distinct color) and reconstructed using the AP SharpTrack toolbox onto coronal diagrams from the Allen Institute Common Coordinate Framework. Fiber tips were located within or just above the caudal portion of the lateral amygdala (black outline), approximately 1.32 to 1.68 caudal to Bregma. Recordings were included if the fiber tip was located within 300 µm of the LA boundary. Scale bar, 1mm.

**Supplemental Figure 2.**
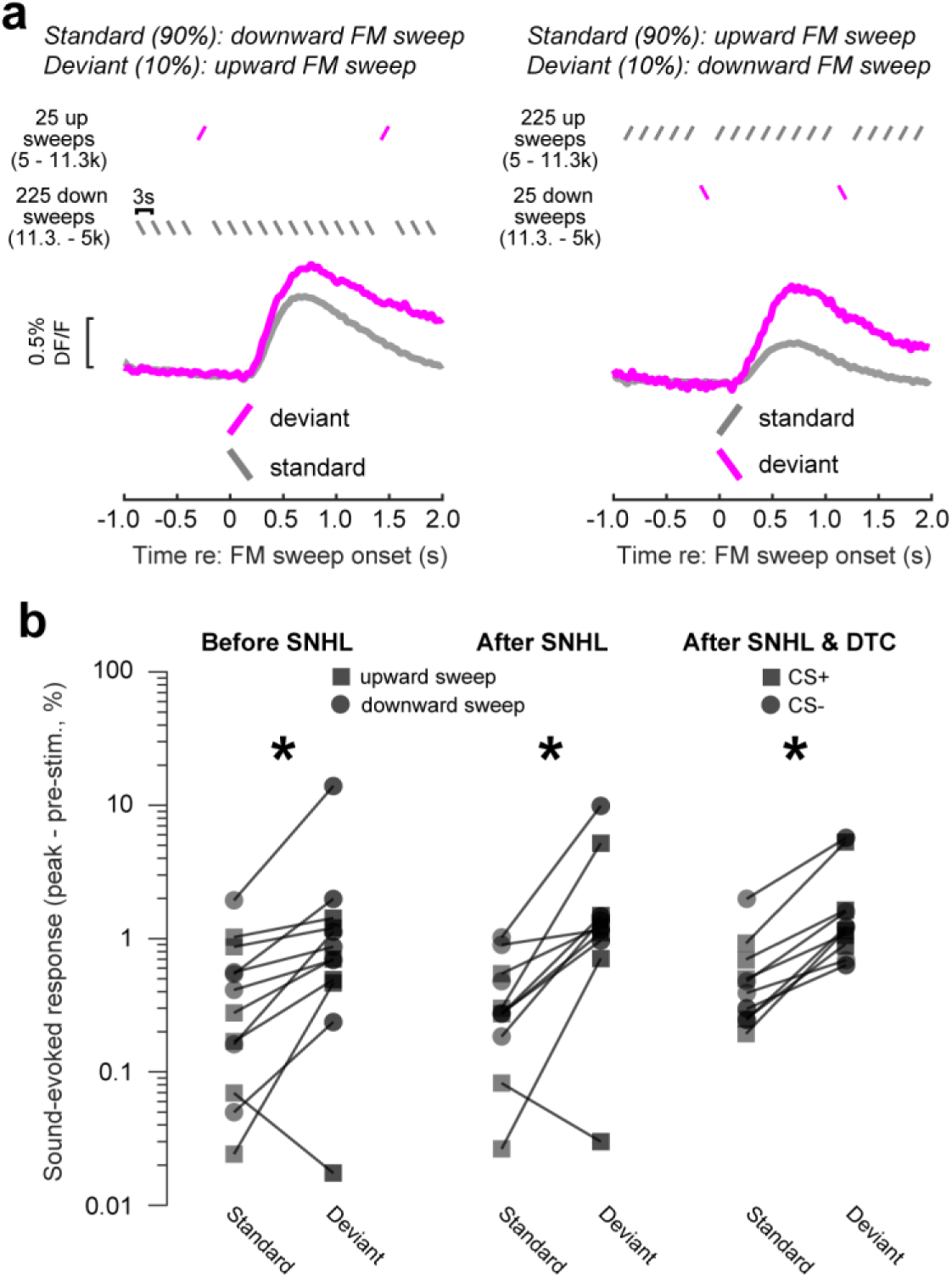
Sensory attributes of CS+ and CS- stimuli are discriminable after NIHL. (a) Oddball paradigm. *Top:* 250 upward (5–11.3 kHz) and downward (11.3–5 kHz) FM sweeps were presented with one direction as the standard (90% of stimuli) and the other as the deviant (10% or stimuli). *Bottom*: Example LA GCaMP responses (ΔF/F, %) to standard (gray) and deviant (magenta) FM sweeps in both standard-deviant configurations. Scale bar, 0.5% ΔF/F. (b) Sound-evoked LA response (peak − pre-stimulus, %) to standard and deviant FM sweeps before NIHL, after NIHL, and after NIHL and DTC (N = 6 PVN-mCherry mice). Squares, CS+; circles, CS−. Deviant responses were significantly greater than standard before NIHL (p = 0.0028) and after NIHL (p = 0.001), and after NIHL & DTC (p = 5.72e-07).

## METHODS

### ANIMAL SUBJECTS

All experimental procedures were approved by the Massachusetts Eye and Ear Institutional Animal Care and Use Committee and conducted in accordance with National Institutes of Health guidelines for the humane care and use of laboratory animals. Data were collected from 34 male and female mice aged 8–9 weeks at the start of recordings. Mice were maintained on a reverse 12 hr light/dark cycle and were provided with *ad libitum* access to food and water.

Fifteen wildtype offspring of hemizygous RBP4-Cre × C57BL/6J breeders contributed to fiber photometry, pupillometry, and discriminative threat conditioning experiments. This genetic cross was selected because they do not exhibit the precocious high-frequency hearing loss observed in C57BL6 mice. Six sham mice participated in all experimental procedures. Nine NIHL mice were used in total: six contributed to passive listening and auditory brainstem response experiments and six contributed to discriminative threat conditioning experiments, with three NIHL mice shared across both experimental series (N = 6/6, sham/NIHL for each experimental series). One NIHL animal was excluded from pupillometry analyses due to an occlusion that prevented reliable pupil tracking. Nineteen PV-Cre mice bred in-house contributed to optogenetic experiments, of whom 11 (N = 6/5, PVN-mCherry/PVN-Chrimson) were used for experiments combining optogenetic manipulation of ACtx PVNs with LA fiber photometry, pupillometry, and discriminative threat conditioning, and 8 (N = 4/2/2, 40-Hz PVN optogenetic/18-Hz PVN optogenetic/40-Hz acoustic stimulation) were used for HO-AC electrophysiology experiments.

### Surgical preparation

#### General methods

Mice were anesthetized with isoflurane in oxygen (5% induction, 0.75–1.5% maintenance). A homeothermic blanket system was used to maintain body temperature at 36.6°C (FHC). Lidocaine hydrochloride was administered subcutaneously to numb the scalp. The dorsal surface of the scalp was retracted and the underlying periosteum removed. The skull surface was cleaned with 70% ethanol.

Virus injections were performed using pulled glass micropipettes (Wiretrol II, Drummond) backfilled with virus solution and delivered at 1 nL/s using a precision injection system (Nanoject III, Drummond). At the conclusion of all surgeries, a titanium headplate (iMaterialise) was affixed to the dorsal surface of the skull with dental cement (C&B Metabond), Buprenex (0.5 mg/kg) and meloxicam (0.1 mg/kg) were administered, and the animal was transferred to a warmed recovery chamber.

#### LA virus injections and optic fiber implantation

For LA injections, we first leveled the head by ensuring that the left and right z coordinates for the lateral skull were within +/- 0.05 mm and the z coordinate of lambda was within +/- 0.05 mm of bregma. A volume of 150-200 nL of AAV9-CaMKII-GCaMP6s was injected (AP: −1.75 mm, ML: +3.45 mm, DV: −3.75 mm relative to bregma) at 1nL/s with a 3–5 s delay between each pulse. The pipette was left in place for 15 minutes following the final pulse before withdrawal. A flat fiber optic cannula (Doric, NA 0.37, 5 mm length, 0.2 mm core diameter, zirconia ferrule) was subsequently implanted in the right hemisphere such that the distal tip terminated 0.2 mm above the injection site. Fibers were secured and optically sealed by applying dental cement mixed with Black India Ink to the exposed skull and head plate. We allowed 3 weeks for viral expression before conducting bulk fiber measurements.

#### Cortical optogenetic experiments

Cortical virus injections were performed during LA injection and fiber implant surgery. The head was tilted 10° such that the pipette tip positioned 2.9mm posterior to bregma passed through the cortical mantle from the primary auditory cortex to more lateral higher-order fields. AAV9-EF1a-DIO-ChrimsonR-mRuby2-KV2.1 or pAAV-hSyn-DIO-mCherry was injected along this trajectory at three points (1000, 750, and 350 µm below the insertion point), delivering 100-150 nL boluses at 1nL/s. At least 10 minutes elapsed between depths before advancing the pipette, and the pipette was left in place for 15 minutes following the final depth before withdrawal. Burr holes were sealed with dental cement following injection. We performed bilateral injections for chronic optogenetic experiments and unilateral (right hemisphere) injections for acute electrophysiology experiments.

#### Chronic bilateral optogenetic stimulation experiments

A small region of the skull overlying the AC of each hemisphere was optically cleared. This was accomplished by applying one layer of each in the following order: cyanoacrylate (Krazy Glue), clear dental cement (C&B metabond), and finally clear nail polish (L.A. colors).

While the nail polish was drying, we affixed cylindrical opaque mating sleeves for optic fiber assemblies (Doric) atop the cleared skull so that light could efficiently reach the ACtx without implanting a fiber on or in the brain. Once all elements had dried, a headplate was affixed to the skull following the procedure described above.

### Noise exposure

Noise exposures were conducted at 8-10 weeks of age during the morning hours. For NIHL and sham exposure, octave-band noise at 16-32 kHz was presented for 2 hours at 103 and 70 dB SPL, respectively. Exposure stimuli were delivered through a high-frequency tweeter (Fostex USA) mounted inside a custom acoustic enclosure (51 × 51 × 51 cm). To minimize standing waves, irregular surface depths and sharp edges were affixed to 3 of the 6 walls using stackable ABS plastic blocks (LEGO) to diffuse the high-frequency sound field. Prior to exposure, mice were placed in a wire-mesh chamber (15 x 15 x 10 cm). This chamber was placed at the center of a continuously rotating plate, ensuring mice were exposed to a uniform sound field.

## DATA ACQUISITION

### Auditory brainstem response measurements

Animals were anesthetized with ketamine (120 mg/kg) and xylazine (12 mg/kg), with supplemental ketamine (60 mg/kg) administered as needed. Body temperature was maintained throughout with a homeothermic heating blanket. Subdermal needle electrodes were placed at the ipsilateral and contralateral pinnae near the tragus, with a ground electrode at the base of the tail. Stimuli were 5 ms tone pips at 8, 11.3, 16, 22.6, and 32 kHz with 0.5 ms rise-fall times, presented at 27 Hz in alternating polarity. Responses were averaged over 512 repetitions per stimulus level, which was incremented in 5 dB steps from 20 to 100 dB SPL. ABR threshold was defined as the lowest stimulus level at which a repeatable waveform could be identified. ABR testing was performed at least 24 days following noise or sham exposure.

### LA fiber imaging and pupillometry experiments

#### Fiber photometry

Bulk GCaMP6s fluorescence was recorded using a dual-wavelength lock-in amplification system, as previously described^26,65^. A 465 nm LED (calcium-dependent signal) and a 405 nm LED (isosbestic control) were sinusoidally modulated at 210 Hz and 330 Hz, respectively, and combined with a fluorescence mini-cube (FMC4, Doric). The optical patch cable was connected to the fiber implant via a zirconia mating sleeve to produce a tip power of 0.1 - 0.2mW. Bulk fluorescent signals were acquired with a femtowatt photoreceiver (2151, Newport) and digital signal processor (Tucker-Davis Technologies RZ5D). The signal was demodulated by the lock-in amplifier implemented in the processor, sampled at 1017Hz and low-pass filtered with a corner frequency at 20Hz. To reduce photobleaching during recordings, fibers were pre-exposed overnight to continuous low-power (<50 µW). Head-fixed recordings were performed double-walled acoustic chamber.

#### Body movements

Mice rested atop an acrylic plate mounted on three piezoelectric force transducers (PCB Piezotronics) coupled to a summation amplifier, allowing measurement of the downward force caused by movement. Force plate signals were digitized at 500 Hz and stored continuously for offline analysis.

#### Pupillometry

Pupil images were captured at 30 Hz with a CMOS camera (Teledyne Dalsa, M2020) equipped with a zoom lens (Tamron 032938) and an infrared longpass filter (Midopt LP830, 25.5 nm cutoff), positioned 25 cm from the right eye. Infrared LEDs (850 nm, Vishay Semiconductors VSLY5850) provided illumination, and ambient visible light was adjusted to maintain an intermediate resting pupil diameter. Pupillometry was performed concurrently with LA fiber photometry across all passive listening and discriminative threat conditioning sessions.

#### Acoustic stimuli

All acoustic stimuli were delivered through a free-field tweeter positioned 20cm from the left ear and calibrated with a ¼” prepolarized microphone (PCB Electronics).

For sensory characterization experiments, we presented pure tones at 8, 11.3, 16, 22.6, and 32 kHz were presented in a pseudorandomized order across 0–80 dB SPL in 20 dB steps (50 ms duration, 15 repetitions per frequency-level combination, 450ms inter-stimulus interval). For LA fiber photometry experiments with concurrent optogenetic stimulation, 8 kHz tones (50 ms duration) and FM sweep trains (5 sweeps, at 1Hz with a 50% duty cycle) presented at 60 dB SPL. Upward (5 to 11.3kHz) and downward (11.3 to 5 kHz) FM sweeps were 0.5s each with 50ms raised cosine onset and offset gating applied at the FM endpoints, producing a sweep velocity of 2.95 octaves/second. To assess stimulus-specific adaptation (SSA), FM sweeps were presented in an oddball configuration during passive listening sessions. The frequent (standard, 90%) and rare (deviant, 10%) stimuli were presented at a 3s onset-to-onset interval (130ms duration). Both FM sweep directions served as standard and deviant in separate configurations (upward sweep as standard/downward sweep as deviant, and vice versa). Each configuration consisted of 250 total stimuli.

DTC Habituation, Recall, and Extinction sessions presented 15 upward and 15 downward FM sweep trains, each 30s in total duration presented at 75 dB SPL in a pseudorandomized order. FM sweeps were 0.5s in duration followed by 0.5s of silence (i.e. a 1 Hz presentation rate), with the same sweep velocity and frequency range described above. The interval between sweep trains specified a 20–180 s inter-trial interval drawn from a decaying exponential distribution to produce a flat hazard function. However, the onset of each sweep train was gated by a real-time motion threshold to avoid instances of complete immobility prior to stimulus onset. This was accomplished by exponentially smoothing the force plate signal and initiating the trial only if supra-threshold movement was for >5 consecutive seconds. To keep each session to approximately 1 hour and minimize photobleaching, a minimum of 8 trials per CS was required for inclusion in the analysis, with 15 trials per CS as the default target.

### Cortical electrophysiology experiments

#### Preparation for acute head-fixed recordings

On the day of recording, mice were briefly anesthetized with isoflurane in oxygen (5% induction, 2% maintenance) and a craniotomy (1.5 × 2 mm) was made around the injection sites, to expose the right auditory cortex. UV-cured composite (Flow-It ALC) was used to construct a recording chamber around the craniotomy, which was filled with optically transparent lubricating ointment (Paralube Vet Ointment) to prevent tissue desiccation. Isoflurane was then discontinued and the mouse was transferred to a dimly lit, double-walled acoustic chamber where it sat head-fixed in a body cradle. Recordings began at least 30 minutes after cessation of anesthesia, as confirmed by normal whisking and grooming behavior.

A 64-channel silicon probe (H3, Cambridge Neurotech) was positioned with a micromanipulator (Narishige) and advanced via a hydraulic microdrive (FHC) into HO-AC at a 15–20° angle to span the non-primary auditory cortex, to a final depth of 1.2–1.6 mm below the pial surface. The brain was allowed to settle for at least 15 minutes before recordings began. Raw neural data were digitized at 32-bit, 24.4 kHz and stored in binary format (PZ5 Neurodigitizer, RZ2 BioAmp Processor, RS4 Data Streamer; Tucker-Davis Technologies). The common-mode signal was subtracted from all channels to eliminate recording artifacts. Signals were notch filtered at 60 Hz and band-pass filtered at 300–3000 Hz (second-order Butterworth filter) for spike detection. Spikes were sorted into single-unit clusters using Kilosort 2.0. Single-unit isolation was confirmed by a refractory period in the interspike interval histogram and an isolation distance greater than 10. Units were classified as regular spiking (RS; putative pyramidal neurons) or fast spiking (FS; putative PV interneurons) based on peak-to-trough delay of the mean spike waveform, with thresholds of >0.55 ms and <0.45 ms, respectively.

#### Acoustic Stimuli

For HO-AC electrophysiology experiments, tonal rate-level functions used tone pips at BF ± 0.5 octaves (100 ms duration, 0–80 dB SPL in 10 dB steps, 20 trials per condition). FM sweep rate-level functions used sweeps with spectral content determined from the frequency response area measured in the same session (500 ms duration, 0–80 dB SPL in 10 dB steps, 500 ms inter-sweep interval, 20 trials per condition). Acoustic stimulation at 40 Hz featured a 33.3 minutes click train (1 μs click duration) at 70 dB SPL.

### Optogenetic stimulation

Light was delivered to the auditory cortex via an optic fiber/ferrule assembly (0.2 mm diameter, 0.22 NA Doric) coupled to a 633 nm diode laser (Omnicron LuxX). The light path was split to provide a light source for each hemisphere separately. Laser power was calibrated for every fiber. For chronic bilateral stimulation experiments through the cleared skull, an opaque mating sleeve (Doric) was attached to each laser patch cable, which fit snugly into the skull-mounted plastic casings. Based on *ex vivo* measurements, the effective power through the cleared skull was found to be approximately 20% of the input power.

For LA fiber photometry experiments with concurrent optogenetic stimulation, light was delivered bilaterally at varying fiber tip power levels, from 0 (off) to 20 mW in a pseudorandom order. For trials employing pure tone stimulation, a 250 ms laser pulse preceding sound onset by 100 ms. For trials presenting longer FM sweep trains, the 5.2 s laser period preceded sound onset by 100ms and consisted of 25 ms pulses were presented at 50% duty cycle (i.e., 20Hz) shaped with a tapering amplitude envelope (0.1s) to reduce rebound excitation.

For cortical electrophysiology experiments, light was delivered through an optic fiber positioned 1–2 mm above the exposed brain surface at 10mW fiber tip power. Continuous stimulation protocols features 33.3 minutes of 40-Hz laser pulses (1 ms pulse width) or 18-Hz laser pulses (2.2 ms pulse width to maintain an 4% duty cycle and equivalent total light energy with 40 Hz stimulation). PVN-mediated inhibition of RS unit spontaneous spiking was assessed with 100ms laser pulses at fiber tip powers ranging from 0 (off) to 20mW, presented in a pseudorandom order (20 repetitions per power level).

For experiments employing 40-Hz optogenetic stimulation followed by chronic LA fiber imaging, mice were head-fixed and received 2000 seconds (∼33.3 minutes) of bilateral stimulation with 1 ms pulse widths, approximately 4 hours before recording or conditioning onset.

### Discriminative threat conditioning

Conditioning sessions were conducted in a different room using a behavioral chamber with a grid floor (Coulbourn Instruments, H10-11M-TC-SF) through which mild foot shocks were delivered (1 s, 0.5 mA AC; Coulbourn Precision Animal Shocker). Stimuli consisted of 5 s FM sweep trains (0.5 s duration, 75 dB SPL, 50 ms raised cosine onset and offset ramps, 1 Hz), during which the fifth sweep of one direction (CS+) co-terminated with the foot shock, while the other direction served as the CS−. CS+/CS− identity was counterbalanced across animals. The apparatus was cleaned with 70% ethanol between sessions.

### Histology

Mice were transcardially perfused with PBS followed by 4% paraformaldehyde. Brains were post-fixed overnight, cryoprotected in 30% sucrose, and sectioned coronally at 50 µm on a cryostat. Sections were mounted and coverslipped with DAPI-containing mounting medium (Fluoroshield, Sigma). GCaMP6s expression and the fiber optic cannula track were visualized using native fluorescence on a widefield epifluorescence microscope.

## DATA ANALYSIS AND STATISTICS

### Fiber photometry analysis

#### Preprocessing

Single-timepoint artifacts exceeding 10 times the median absolute deviation of the raw signal were identified and removed prior to further processing. Fluorescence signals were converted to ΔF/F by calculating slow non-specific changes in fluorescence (F₀) with a 60-second sliding median filter, subtracting F₀ from the raw signal, applying a 3-sample median filter and dividing by F₀. To reduce potential contributions of intrinsic signals and movement artifacts, analyses were performed on a corrected GCaMP signal in which the fractional change in fluorescence measured with the 405nm excitation was smoothed using a 1 s gaussian filter and then subtracted from the 465nm signal for each trial.

#### Stimulus-evoked responses

Sound-evoked LA responses were quantified from trial-averaged ΔF/F traces as the mean ΔF/F within a post-stimulus response window. For sensory characterization experiments, the response at each sound level was quantified as the mean ΔF/F within a 0.05–1.0 s post-stimulus window, following subtraction of the mean pre-stimulus baseline (1.0 s prior to sound onset). To compute a single response value across the sound level growth function, we calculated the area under the curve using trapezoidal integration across 40, 60, and 80 dB SPL (‘trapz’, Matlab). Changes in responsivity across sessions were quantified with an evoked response asymmetry index, defined as ([Session X – Baseline_mean_] / [Session X + Baseline_mean_]), where Baseline_mean_ is the mean AUC across the two pre-exposure sessions.

For experiments that measured real-time modulation of LA responses during optogenetic activation, the laser onset preceded sound onset by 100ms. The response window spanned 0.2–1.5 s post-stimulus onset for the 50ms tone burst and 0.2–5.2 s for the 1Hz FM sweep train. A suppression asymmetry index quantified laser-induced changes independently for each session as (AUC_PowerX – AUC_Off) / (|AUC_PowerX| + |AUC_Off|), where the reference is the off (0 mW) condition. To compare changes with laser stimulation to responses at baseline, the asymmetry index was calculated as (AUC_Power X – AUC_Off_Baseline_) / (|AUC_Power X| + |AUC_Off_Baseline_|). Absolute values were used in the denominator because cortical PVN activation could suppress LA responses below zero. For oddball sessions (SSA), the peak ΔF/F within a 1-s post-stimulus window was taken as the response amplitude for standard and deviant trials separately.

For the analysis of CS+ and CS- responses from DTC sessions, the positive area under the curve from the trial-averaged response was computed with the ‘trapz’ function in Matlab across the full 30-s CS period. A CS asymmetry index was computed as (AUC_CS+ − AUC_CS−) / (AUC_CS+ + AUC_CS−). To compare response magnitudes across sessions, AUC values were expressed as fold change relative to the mean AUC across the two habituation sessions.

### Electrophysiology analysis

Sound-responsive RS units were identified by a significant difference in pre- vs post-stimulus spike rates across a 30-80 dB SPL range (paired t-test, p < 0.05). For tone-evoked sound level growth functions, evoked spike rates 100 ms post-stimulus onset were averaged across three frequencies (BF ± 0.5 octaves) at each sound level. For FM sweeps, the -100 – 0ms pre-stimulus spike rate was subtracted from spike rate measured 0 - 500 ms post-stimulus onset. Each unit’s rate-level function was normalized to its maximum firing rate at baseline.

For PVN-mediated feedforward inhibition, RS unit spike rates were measured during a 150 ms window beginning 5 ms after laser onset. Only units with a minimum spontaneous firing rate of ≥ 0.5 Hz were included. A PVN inhibition asymmetry index was computed as (FR_XmW − FR_0mW) / (FR_XmW + FR_0mW), where FR is the mean spike rate during the response window and the 0 mW (laser off) condition serves as the reference. Units were included only if data were available across all three timepoints.

### Pupillometry analysis

Pupil diameter was estimated from eight DeepLabCut-tracked landmarks placed at the cardinal and intercardinal positions along the pupil margin. A ResNet-50-based neural network was trained on 95% of labeled frames using default DeepLabCut parameters. Tracked landmark coordinates were used to fit an ellipse to the pupil boundary using a least-squares criterion; the long axis of the fitted ellipse was taken as pupil diameter for each frame. Frames in which more than 25% of markers fell below a likelihood threshold of 0.8 were discarded.

Pupil traces were resampled to a common 30 Hz time base spanning −1 to +4 s relative to stimulus onset. The sound-evoked pupil response was expressed as a fractional change in pupil diameter (ΔP/P₀), where P₀ is the mean pupil diameter during the 1-s pre-stimulus baseline, computed separately for each trial.

Sound-evoked pupil responses were quantified from ΔP/P₀ traces. The mean ΔP/P₀ within a 2-s window beginning at stimulus onset was taken as the evoked response amplitude for each trial. For each animal and session, evoked responses were averaged across trials at each sound level, and the area under the pupil rate-level function was computed by trapezoidal integration across a 40 - 80 dB SPL range.

#### LA–pupil coupling analysis

Trial-by-trial temporal coupling between LA GCaMP and pupil signals was quantified using normalized cross-correlation. For each trial, the LA ΔF/F trace within a 2-s post-stimulus window was resampled from 1017 Hz to 30 Hz to match the pupil sampling rate, and the pupil ΔP/P₀ trace was smoothed with a 5-sample moving average. Normalized cross-correlations were computed with a maximum lag of ±2 s and averaged across trials within each animal. Coupling strength was defined as the integral of the positive portion of the mean cross-correlogram.

### Freezing

Freezing was quantified from piezoelectric force plate signals sampled at 500 Hz during Habituation, Recall, and Extinction sessions^66–68^. The force signal was divided into sliding 1-s windows (0.5-s step) and the power spectrum was computed for each window via Fast Fourier Transform (FFT). Mean spectral power at ≤ 5 Hz was used as the movement metric. Windows in which low-frequency power fell below threshold were classified as freezing; thresholds were calibrated per animal to account for variation in posture and plate coupling. Freezing bouts shorter than 2 s were excluded. Freezing was scored during the 30-s CS presentation and expressed as the percentage of time spent freezing. A freezing CS asymmetry index was computed as (Freeze_CS+ − Freeze_CS−)/ (Freeze_CS+ + Freeze_CS−).

### Post-mortem fiber tip reconstructions

Fiber tip localization was performed using the AP_histology pipeline (Bhagat & Bhagat; github.com/petersaj/AP_histology), an updated implementation of the SHARP-Track framework^69^, registered to the Allen Mouse Brain Common Coordinate Framework (CCF v3). The section containing the center of the fiber tip was identified to calculate the dorsal-ventral distance between the fiber tip and LA boundary. A distance < 300 μm was used to ensure that cone of light from the fiber tip would include GCaMP from the LA.

### Statistical analysis

All statistical analyses were performed in MATLAB 2023b (MathWorks). Group comparisons were performed using linear mixed-effects models (LMEs) with mouse (or unit, for electrophysiology analyses) as a random intercept. Freezing rates during CS+ and CS− presentations were compared within groups using paired t-tests. Post hoc pairwise comparisons were corrected for multiple comparisons using the Holm–Bonferroni procedure. Statistical results are provided in the figure legends. Unless otherwise stated, error shown is the standard error of the mean (SEM).

## ACKNOWLEDGEMENTS

This work was supported by NIH grants DC017078 (DBP), DC009836 (DBP), DC022957 (BA), a research grant from the Nancy Lurie Marks Family Foundation (DBP), a research grant from the Misophonia Research Fund (BA), and a hyperacusis research grant from the Hearing Health Foundation (BA).

We are grateful to Molly Persky for support with mouse colony maintenance, Jennifer Zhu for post-mortem tissue processing, and Ethan Lawler for supporting analysis of the post-mortem tissue.

